# Copper Induces Zebrafish Central Neural System Myelin Defects: the Regulatory Mechanisms in Wnt/Notch-*hoxb5b* Signaling and Underlying DNA Methylation

**DOI:** 10.1101/2019.12.16.877860

**Authors:** Ting Zhang, PengPeng Guan, Guang Zhao, YaPing Fang, Hui Fu, Jian-Fang Gui, GuoLiang Li, Jing-Xia Liu

## Abstract

Unbalanced copper (Cu^2+^) homeostasis is associated with neurological development defects and diseases. However, the molecular mechanisms remain elusive. Here, central neural system (CNS) myelin defects and down-regulated expression of Wnt/Notch signaling and their down-stream mediator *hoxb5b* were observed in Cu^2+^ stressed zebrafish larvae. Loss/knockdown-of-function of *hoxb5b* phenocopied the myelin and axon defects observed in Cu^2+^ stressed embryos. Meanwhile, activation of Wnt/Notch signaling and ectopic expression of *hoxb5b* could rescue copper-induced myelin defects, suggesting Wnt&Notch-*hoxb5b* axis mediated Cu^2+^ induced myelin and axon defects. Additionally, whole genome DNA methylation sequencing unveiled that a novel gene *fam168b*, similar to *pou3f1*/*2*, exhibited significant promoter hypermethylation and reduced expression in Cu^2+^ stressed embryos. The hypermethylated locus in *fam168b* promoter acted pivotally in its transcription, and loss/knockdown of *fam168b*/*pou3f1* also induced myelin defects. Moreover, this study unveiled that *fam168b*/*pou3f1* and *hoxb5b* axis acted in a seesaw manner during fish embryogenesis, and demonstrated that copper induced the down-regulated expression of the Wnt&Notch-*hoxb5b* axis dependent of the function of copper transporter *cox17*, coupled with the promoter methylation of genes *fam168b/pou3f1* and their subsequent down-regulated expression dependent of the function of another transporter *atp7b*, making joint contributions to myelin defects in embryos. Those data will shed some light on the linkage of unbalanced copper homeostasis with specific gene promoter methylation and signaling transduction as well as the resultant neurological development defects and diseases.

**Author summary:** In this study, we first unveiled that copper induced central neural system (CNS) myelin defects *via* down-regulating Wnt/Notch-*hoxb5b* signaling, and parallel with hypermethylating promoters of genes *fam168b*/*pou3f2* and their subsequent down-regulated expression. Additionally, we unveiled that *fam168b*/*pou3f1* and *hoxb5b* axis acted in a seesaw manner during fish embryogenesis. Genetically, we unveiled that copper was trafficked to mitochondrion *via cox17* then led to the down-regulation of Wnt&Notch-*hoxb5b* axis, and was trafficked to trans-Golgi network *via atp7b* to induce the hypermethylation and the down-regulated expression of *pou3f1*/*fam168b* genes, making joint contributions to myelin defects in embryos.

## Introduction

Many neurological diseases with behavioral changes and neurological disorders are associated with the unbalanced copper homeostasis in human, such as Alzheimer’s disease (AD), Wilson’s disease (WD) and Wallerian degeneration (WD)(1, 2). Excess copper has been reported to damage nervous system and lead to the behavioral abnormalities in fish (3, 4). However, the detailed molecular characteristics and the potential mechanisms underlying copper-induced neural defects, especially axonal and myelin defects, remain unclear.

Axons transmit electrical impulses to neuron’s targets, which is an essential process for the establishment of the nervous system. Axonal damage has been shown to cause neurological disorders, such as stroke, traumatic brain/spinal cord injuries, multiple sclerosis (MS) and Wallerian degeneration (WD) (5, 6). Myelin sheaths are essential for the rapid and efficient propagation of action potentials as well as for the support for the integrity of axons in the vertebrate nervous system. In the central nervous system (CNS), oligodendrocytes spirally wrap axons in multilamellar plasma membrane and eventually compact to form the myelin sheaths (7, 8). The compacted myelin sheaths increase the resistance of axons and reduce their capacitance by several orders of magnitude(8, 9). The failure of compacted myelin formation leads to delayed or interrupted signal conduction, contributing to motor, sensory, and cognitive behavioral deficits(10, 11). Even a subtle defect of CNS myelin can cause a persistent cortical network dysfunction and induce neuropsychiatric disorders in mouse(12–14). Myelin disorder in human has been reported to associate with a series of neurodegenerative diseases such as MS, Menkes diseases (MD), Parkinson’s diseases (PD) and Huntington’s diseases (HD) (14, 15).

DNA methylation is implicated in many copper-induced disorders, including AD, WD, MS and MD (16, 17). In WD patients, accumulated copper dysregulates methylation status (17, 18). The methylation level of PAD2 is reduced in copper-accumulated MS brain, leading to changes of *mbp* expression (19, 20). Copper has been reported to upregulate the expression of DNA methylation- and stress-related genes in zebrafish (21, 22). However, few reports are available about copper-induced locus-specific DNA methylation during embryogenesis, and few studies have linked this methylation with its induced myelin developmental damages in vertebrates.

The normal trafficking of the copper is essential for normal cellular functions. In vertebrate, copper is delivered to different organelle by independent copper chaperones, such as *cox17*, *atp7b*, and etc(23).. *Cox17* helps copper being delivered to mitochondrial cytochrome c oxidase, and Trans-Golgi network (TGN) resident *atp7b* helps copper being pumped to the circulation or to extracellular (24, 25).

In this study, zebrafish *in vivo* system was used to investigate the cellular and molecular mechanisms of copper-induced CNS myelin defects. Copper was revealed to induce myelin defects *via* down-regulating the expression of Wnt&Notch-*hoxb5b* axis. Additionally, the gene *fam168b*, similar to *pou3f1* and *pou3f2*, showed hypermethylation in the promoter regions and reduced expressions in copper stressed embryos, which might mediate in a parallel pathway to Wnt&Notch-*hoxb5b* in copper-induced myelin defects. Moreover, *cox17^-/-^*and *atp7b*^-/-^ mutants were used to verify the correlation of copper trafficking deficiency in specific organelle and the occurrence of developmental myelin defects in copper stressed embryos in this study.

## Results

### Cu^2+^ induces embryonic CNS myelin and axon defects in zebrafish

Transmission electron microscopy (TEM) detection revealed compacted myelin sheaths in the spinal cord in the control larvae at 5 dpf (Fig 1A1-A2), but significantly thinner (increased g-ratio) and uncompacted myelin in the spinal cord in Cu^2+^ stressed larvae (Figs 1A3-A5). Additionally, WISH assays detected significantly reduced expression of *mbp*，*plp1a*, and *olig2* (Figs 1B, S1A, and S1B) in the spinal cord in Cu^2+^ stressed embryos at 96 hpf. qRT-PCR results further conformed the reduced expression of *mbp* and *mpz* in the Cu^2+^ stressed embryos at 96 hpf (Fig 1C). The Cu^2+^ effects on myelin and axon development were further tested by analyzing the fluorescence in Cu^2+^ stressed transgenic *Tg*(*mbp*:EGFP) and *Tg*(*olig2*:dsRED) embryos, with a significant down-regulation observed in their respective fluorescence at 96 and 48 hpf (Fig 1D). Furthermore, the length of axons was remarkably reduced in Cu^2+^ stressed embryos (Fig 1E).

**Fig 1.**
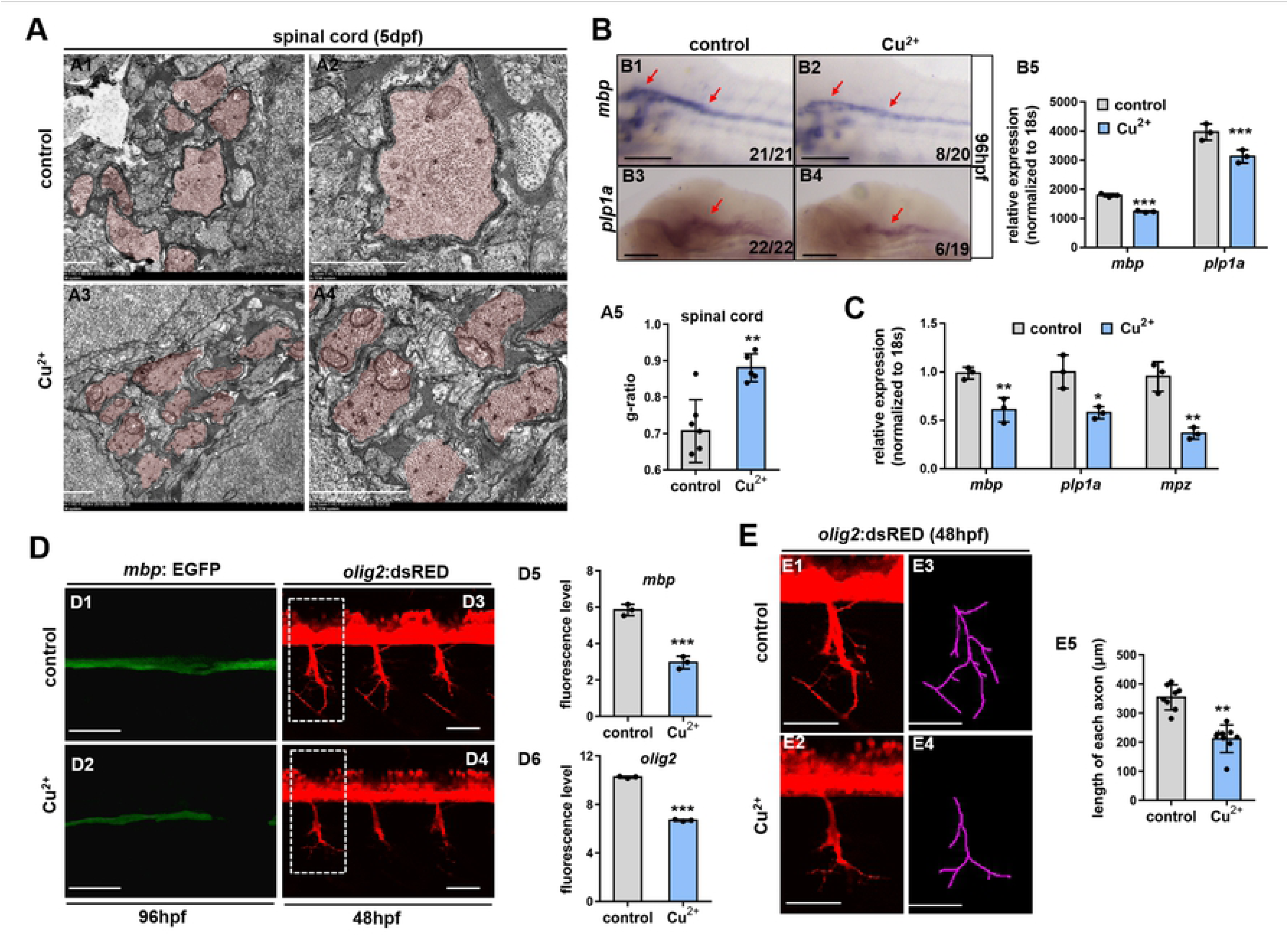
Central neural system (CNS) myelin and axon defects in Cu^2+^ stressed zebrafish embryos. (**A**) Transmission electron micrographs of transverse ventral spinal cord sections in the control or Cu^2+^ stressed larvae at 5dpf (days post fertilization). Myelinated axons are pseudocolored in red (**A1-A4**). Scatter plots of the myelin g-ratios in the control or Cu2+ stressed larvae (**A5**). (**B**) WISH analysis of CNS myelin marker *mbp* expression in the control or Cu^2+^ stressed larvae at 96 hpf (hours post fertilization) (**B1-B2**), and quantification analysis of the WISH data in different samples (**B3**). (**C**) qRT-PCR analysis of the expression in CNS myelin markers *mbp* and *mpz* in the control or Cu^2+^ stressed larvae at 96 hpf. (**D**) Confocal micrographs of 96-hpf *Tg*(*mbp*:EGFP) (**D1-D2**) and 48-hpf *Tg*(*olig2*:dsRED) (**D3-D4**) in the control or Cu^2+^ stressed embryos, and quantification analysis of fluorescence level in different samples (**D5-D6**). (**E**) The length of axon in the control or Cu^2+^ stressed embryos at 48 hpf. The axons were traced (**E3-E4**) and measured (**E5**) by Neuron J software. Each experiment was repeated three times, and a representative result is shown. Data are mean ± SD. **B1-B2**, **D1-D4**, **E1-E2**, lateral view, anterior to the left and dorsal to the up. The red arrow indicates mbp-expression in the spinal cord. \**P* < 0.05, \*\**P* < 0.01, \*\*\**P* < 0.001. Scale bars, 1 μm (A), 100 μm (B) and 50μm (E). See also Fig S1.

### CNS myelin and axon formation in *hoxb5b* loss- and gain-of-function embryos

*Hoxb5b* exhibited significantly and specifically reduced expression in Cu^2+^ stressed embryos (22), and was reported to function importantly in axon guidance in mouse(26). Thus, in this study, *hoxb5b* was assumed to be a potential mediator in Cu^2+^ induced myelin and axon defects. To validate the hypothesis, an anti-sense morpholino (*hoxb5b*-MO) and a *hoxb5b* null mutant with a 4-bp deletion in the first exon (*hoxb5b*^-/-^) (Fig 2A) were applied to test the *hoxb5b* roles in Cu^2+^ induced myelin and axon defects. Embryos injected with *hoxb5b*-MO at 48 hpf exhibited brain hypoplasia, eye hypoplasia, trunk abnormalities, and reduced body size (Fig 2B2), which phenocopied the defects observed in Cu^2+^ stressed embryos. However, *hoxb5b*^-/-^ mutant embryos exhibited almost normal-like phenotype at 48 hpf (Figs 2B3, B4).

**Fig 2.**
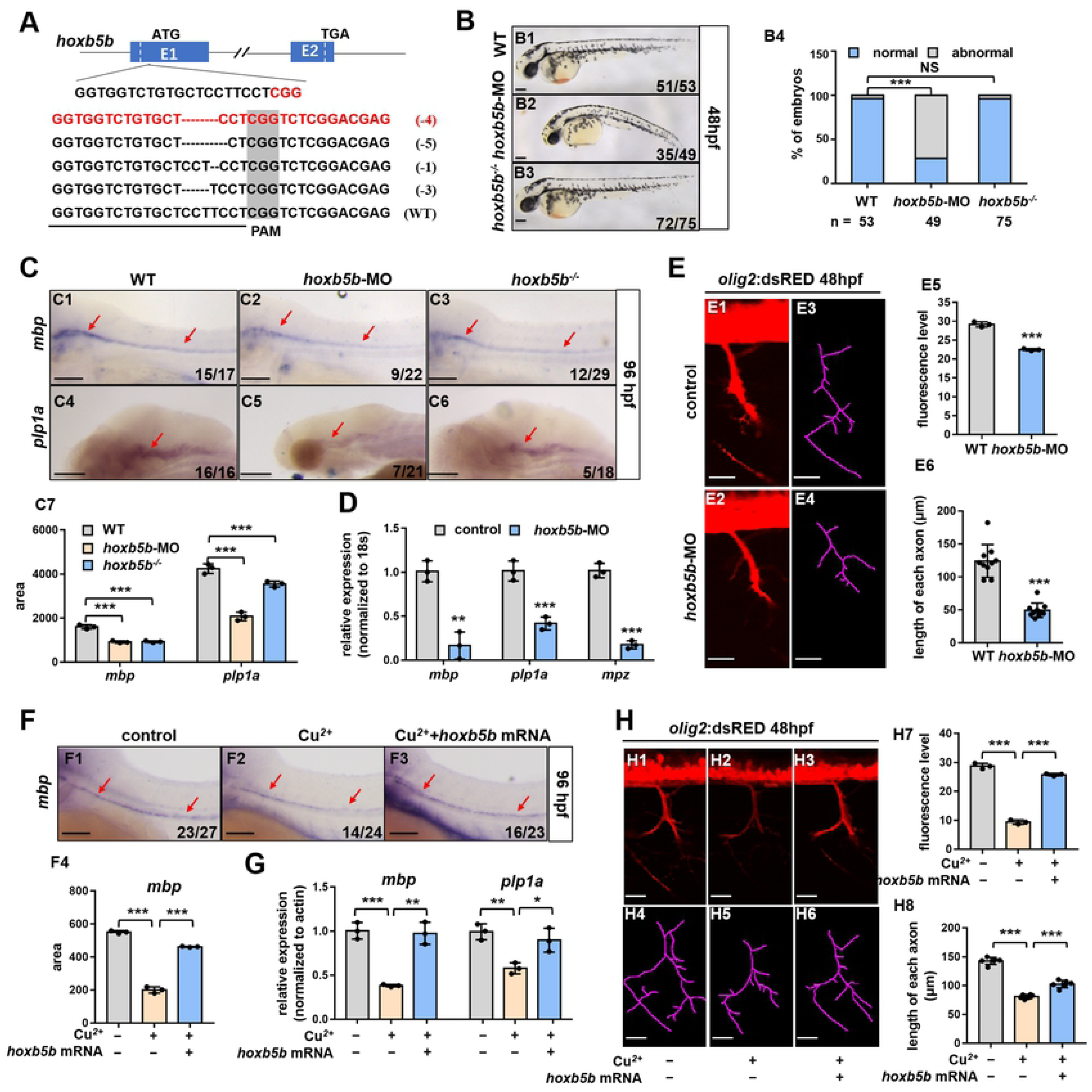
CNS myelin and axon formation in *hoxb5b* loss- and gain-of-function embryos. (**A**) Schematic diagram showing the genomic structure and a genetic mutation of zebrafish *hoxb5b* gene, with the red line indicating the genotypic deletion of the mutation used in this study. ATG denotes the translation start codon; TGA, the translation termination codon; PAM, the protospacer adjacent motif; slash, intron; blue horizontal bar, exon; dotted lines, the deletion of *hoxb5b*; numbers, the length of mutant base. (**B**) Phenotypes of WT embryos (**B1**), WT embryos injected with *hoxb5b*-MO (B2), or *hoxb5b*^-/-^ mutant embryos (**B3**) at 48 hpf, and the percentage of embryos exhibiting abnormal development in different samples (**B4**). (**C**) WISH analysis of CNS myelin marker *mbp* (**C1-C3**) expression in WT embryos, WT embryos injected with *hoxb5b*-MO, or *hoxb5b*^-/-^ mutant embryos at 96 hpf, and quantification analysis of the WISH data in different samples (**C4)**. (**D**) qRT-PCR expression analysis of CNS myelin markers *mbp* and *mpz* in WT embryos, WT embryos injected with *hoxb5b*-MO, or *hoxb5b*^-/-^ mutants at 96 hpf. (**E**) Confocal micrographs of *Tg*(*olig2*:dsRED) in the control or *hoxb5b*-MO injected embryos at 48 hpf (**E1-E2**), and quantification analysis of fluorescence levels (**E5**) in different samples. Tracings (**E3-E4**) and length of axon (**E6**) in different samples. (**F**) WISH analysis of CNS myelin marker *mbp* expression in the control, Cu^2+^ stressed, or Cu^2+^ stressed embryos with ectopic *hoxb5b* expression at 96 hpf (**F1-F3**), and quantification analysis of the WISH data in different samples (**F4**). (**G**) qRT-PCR expression analysis of CNS myelin marker *mbp* in the control, Cu^2+^ stressed, or Cu^2+^ stressed embryos with ectopic *hoxb5b* expression at 96 hpf. (**H**) Confocal micrographs of *Tg*(*olig2*:dsRED) in the control, Cu^2+^ stressed, or Cu^2+^ stressed embryos with ectopic *hoxb5b* expression at 48 hpf (**H1-H3**). Quantification analysis of fluorescence levels in different samples (**H7**). Tracings (**H4-H6**) and length of axon (**H8**) in different samples. Each experiment was repeated three times, and a representative result is shown. Data are mean ± SD. **B1-B3, C1-C3, E1-E2, F1-F3, H1-H3**, lateral view, anterior to the left and dorsal to the up. The red arrow indicates *mbp*-expression in the spinal cord. \**P* < 0.05, \*\**P* < 0.01, \*\*\**P* < 0.001. NS, not significant. Scale bars, 200 μm (B),100 μm (C, F) and 20 μm (E, H). See also Fig S2.

Additionally, compared with the WT larvae, *hoxb5b* morphants or *hoxb5b^-/-^* mutants exhibited significantly decreased expression in the CNS myelin markers *mbp* and *plp1a* at 96 hpf (Figs 2C, 2D S2A1, and S2A2), identical to that in Cu^2+^ stressed larvae. The *olig2* promoter driven fluoresce was observed to be significantly down-regulated in *hoxb5b*-MO injected *olig2*:dsRED transgenic embryos, compared with that in the control embryos at 48 hpf (Fig 2E). Furthermore, the length of each axon was significantly reduced in *hoxb5b* morphants (Fig 2E6), which also phenocopied the defects observed in Cu^2+^ stressed embryos.

The expression of CNS myelin markers *mbp* and *plp1a* (Figs 2F, 2G and S2A3) and the *olig2* promoter driven fluoresce in CNS and the length of axon (Fig 2H) were partially rescued in Cu^2+^ stressed embryos *via* ectopic expression of *hoxb5b*.

### Activation of Wnt or Notch signaling rescues myelin and axon defects in Cu^2+^ stressed embryos *v ia* recovering *hoxb5b* expression

The microarray data showed that the expressions of Wnt and Notch signaling genes were reduced in Cu^2+^ stressed embryos (Figs 3A, S2B1 and Table S9), and qRT-PCR assays confirmed the down-regulated expressions of Wnt signaling(27) and Notch signaling genes (Fig S2B2) in Cu^2+^ stressed embryos. It has been reported that both Wnt and Notch signaling specified the oligodendrocyte fate (1, 28–30), and *hoxb5b*, is downstream of these two signaling pathways(31, 32). In this study, both Wnt agonist BIO and NICD *notch3* mRNA partially rescued *hoxb5b* expression in Cu^2+^ stressed embryos separately (Figs 3B, 3C). WISH and qRT-PCR analysis exhibited the expression of *mbp* and *plp1a* was recovered to nearly normal level in Cu^2+^ stressed embryos co-exposed with BIO (Figs 3D, 3E). Additionally, BIO significantly recovered the down-regulated fluorescence level and the length of the fluorescent axon to nearly normal level in Cu^2+^ stressed *olig2*:dsRED transgenic embryos (Fig 3F).

**Fig 3.**
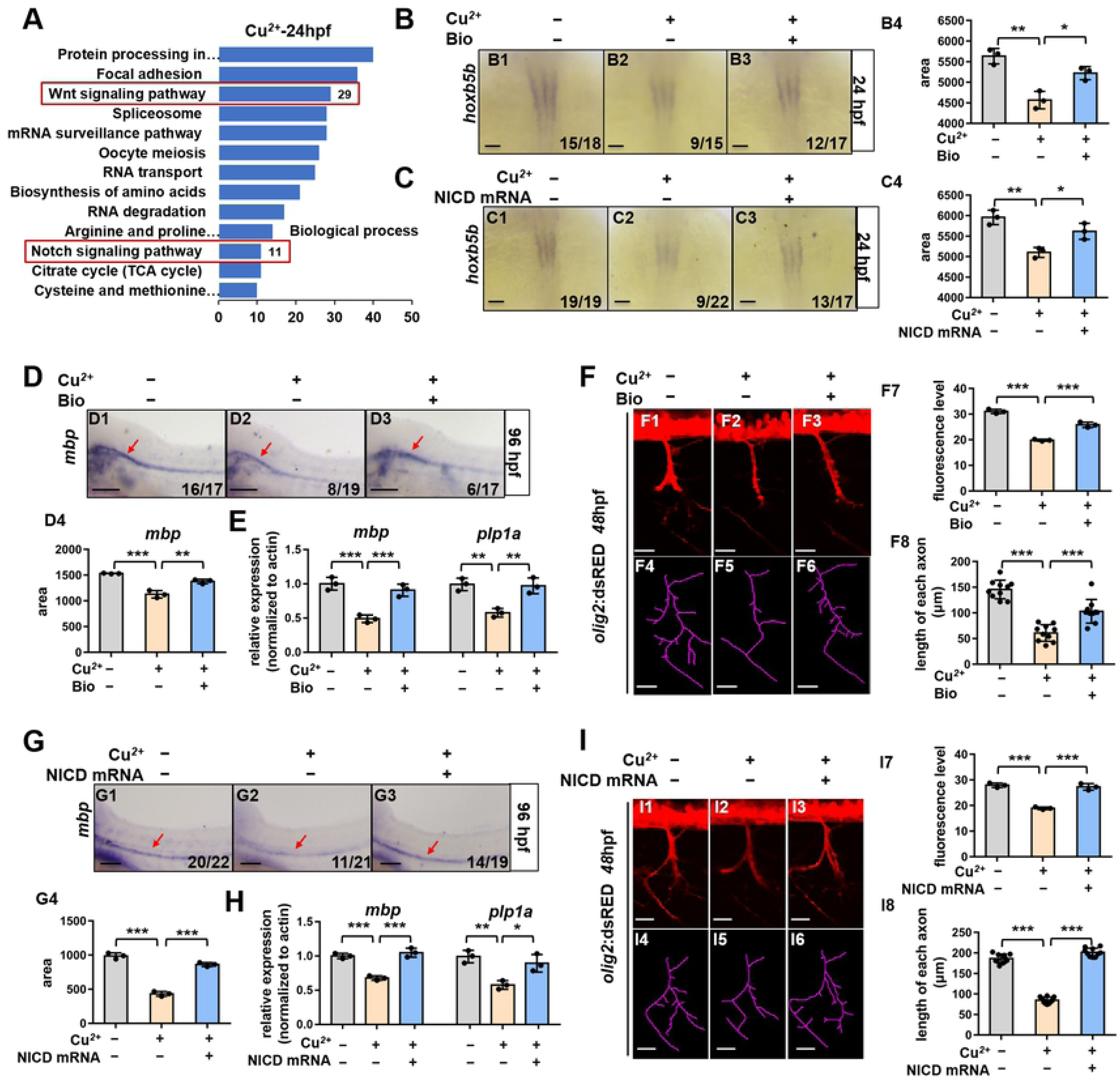
Activation of Wnt or Notch signaling rescues myelin and axon defects in Cu^2+^ stressed embryos via recovering *hoxb5b* expression. (**A**) Enrichment of Wnt and Notch signaling pathways for down-regulated genes in 24 hpf Cu^2+^ stressed embryos *via* KEGG pathway analysis. (**B**) WISH analysis of *hoxb5b* expression in the control, Cu^2+^ exposed, or Cu^2+^ and BIO co-exposed embryos at 24 hpf (**B1-B3**), and quantification analysis of the WISH data in different samples (**B4**). (**C**) WISH analysis of *hoxb5b* expression in the control, Cu^2+^ exposed, or Cu^2+^ and BIO co-exposed embryos with ectopic NICD expression at 24 hpf (**C1-C3**), and quantification analysis of the WISH data in different samples (**C4**). (**D**) WISH analysis of CNS myelin marker *mbp* expression in the control, Cu^2+^ exposed, or Cu^2+^ and BIO co-exposed embryos at 96 hpf (**D1-D3**). Quantification analysis of the WISH data in different samples (**D4**). (E) Confocal micrographs of *Tg*(*olig2*:dsRED) in the control, Cu^2+^ exposed, or Cu^2+^ and BIO co-exposed at 48 hpf (**E1-E3**). Quantification analysis of fluorescence levels (**E7**) in different samples. Tracings (**E4-E6**) and length of axon in different samples (**E8**). (**F**) WISH analysis of CNS myelin marker *mbp* expression in the control, Cu^2+^ exposed, or Cu^2+^ and BIO co-exposed embryos with NICD ectopic expression at 96 hpf (**F1-F3**). Quantification analysis of the WISH data in different samples (**F4**). (**G**) Confocal micrographs of *Tg*(*olig2*:dsRED) in the control, Cu^2+^ exposed or Cu^2+^ and BIO co-exposed embryos with ectopic NICD expression at 48 hpf (**G1-G3**). Quantification analysis of fluorescence levels in different samples (**G7**). Tracings (**G4-G6**) and length of axon in different samples (**G8**). Each experiment was repeated three times, and a representative result is shown. Data are mean ± SD. **B1-B3, C1-C3,** dorsal view, anterior to the top, **D1-D3, E1-E3, F1-F3, G1-G3**, lateral view, anterior to the left and dorsal to the up. The red arrow indicates mbp-expression in the spinal cord. \**P* < 0.05, \*\**P* < 0.01, \*\*\**P* < 0.001. NS, not significant. Scale bars, 200 μm (B,C),100 μm (D, F) and 20 μm (E, G). See also Fig S2.

WISH and qRT-PCR assays showed the expression of *mbp* and *plp1a* was significantly rescued *via* ectopic expression of NICD *notch3* mRNA in Cu^2+^ stressed larvae (Figs 3G, 3H, and S2B3), and the fluorescence for the expression of *olig2* and the length of fluorescent axon was partially rescued in the Cu^2+^ stressed *olig2*:dsRED transgenic embryos with ectopic expression of NICD mRNA (Fig 3I).

### DNA methylation and transcriptional activity of myelin genes in Cu^2+^ stressed embryos

Under stress conditions, epigenetic DNA methylation has been reported to function importantly in disease process and intergenerational inheritance (33, 34). Thus, the whole genome methylation level in Cu^2+^ stressed larvae was examined to unveil the potential epigenetic mechanisms underlying Cu^2+^ induced myelin and axon defects. It has been unveiled that Cu^2+^ induced the expression of 26 hyper-methylated and 31 hypo-methylated genes in Cu^2+^ stressed larvae(35). Among them, genes *pou3f1*, *pou3f2* and *fam168b*, which were associated with myelin and axon, were hypermethylated in the promoter domain (Figs 4A, S3A1 and S3B). Based on the microarray data reported previously(22), this study unveiled the down-regulated expression of genes *pou3f1*, *pou3f2*, and *fam168a*, the homolog gene of *fam168b* in Cu^2+^ stressed embryos (Fig S3A2), which was further confirmed by qRT-PCR analysis (Fig 4B1). Specifically, *pou3f1* showed obviously decreased expression in the brain of Cu^2+^ stressed larvae (Figs 4B2-B4 and S3A3). Additionally, the expression of *fam168b* was also significantly reduced in Cu^2+^ stressed larvae from 24 hpf to 96 hpf (Figs 4B1 and C), and its promoter exhibited significant hypermethylation at 96 hpf (Figs 4D and S3C).

**Fig 4.**
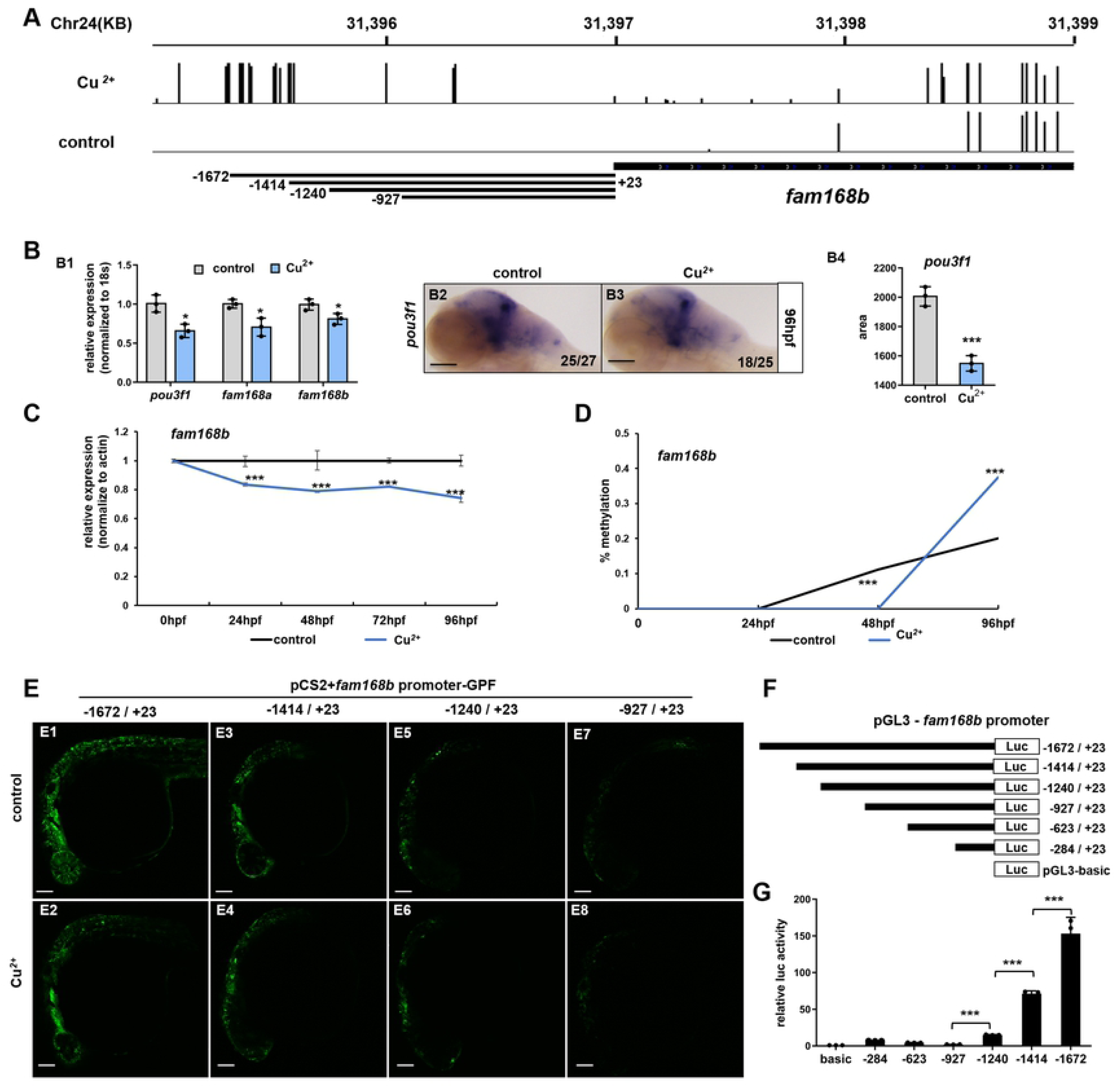
DNA methylation and transcriptional activity of gene *fam168b* in Cu^2+^ stressed embryos. (**A**) Graphical representation of methylation patterns in the promoter domain of *fam168b* gene in the control or Cu^2+^ stressed larvae at 96 hpf. (**B**) qRT-PCR analysis of *pou3f1*, *fam168a*, and *fam168b* expression in the control or Cu^2+^ stressed embryos (**B1**). WISH analysis of expression for the myelination transcriptional factor *pou3f1* in the control or Cu^2+^ stressed larvae at 96 hpf (**B2-B3**), and quantification analysis of the WISH data in different samples (**B4**). (**C**) qRT-PCR analysis of *fam168b* expression in Cu^2+^ stressed embryos at different developmental stages. (**D**) Bisulfite PCR sequencing analysis of *fam168b* methylation in Cu^2+^ stressed embryos at different stages. (**E**) Representative embryos injected with different 5’ unidirectional deletions of *fam168b* promoter driven GFP plasmid at 24 hpf. A series of plasmids containing 5’ unidirectional deletions of *fam168b* promoter region (−1672, −1414, −1240 and −927) were injected separately into zebrafish embryos at one-cell stage, and embryos at 24 hpf were observed *via* confocal microscope. (F) The schematic of truncated *fam168b* promoter mutants. (G) The luciferase activities of different truncated *fam168b* promoters in HEK293 cells. Each experiment was repeated three times, and a representative result is shown. Data are mean ± SD. **B2-B3, E1-E8**, lateral view, anterior to the left and dorsal to the up. \**P* < 0.05, \*\**P* < 0.01, \*\*\**P* < 0.001. NS, not significant. Scale bars, 100 μm. See also Fig S3.

**Fig 5.**
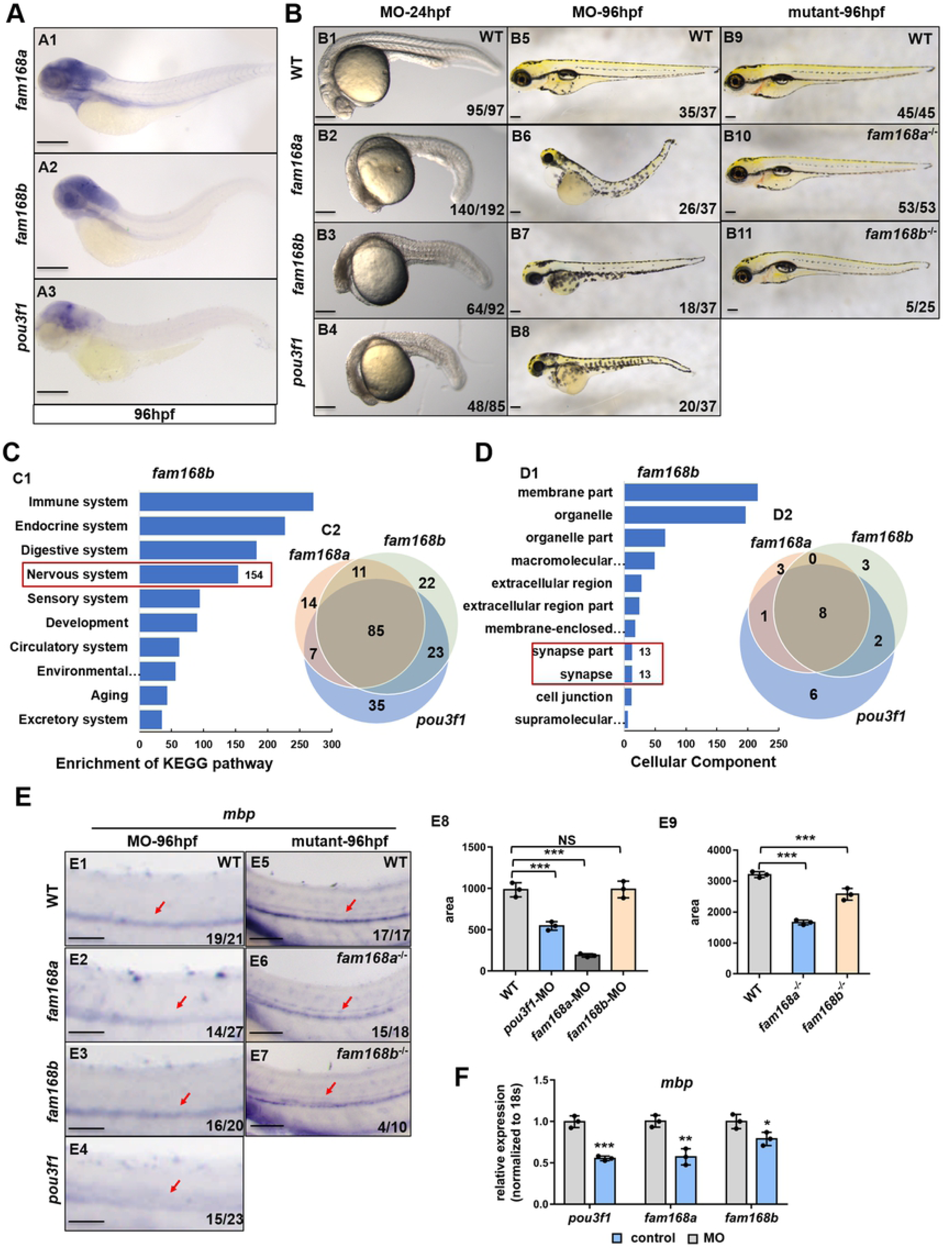
Myelin and axon formation in *fam168a*/*fam168b* loss- and gain-of-function embryos. (**A**) Expression and distribution of *fam168a* and *fam168b* in WT embryos at 96 hpf, which exhibited similar expression pattern to that of myelination transcriptional factor *pou3f1* in embryos. (**B**) Phenotypes of WT embryos, *fam168a*, *fam168b* and *pou3f1* morphants or mutants at 24 hpf and 96 hpf. (**C**) Enrichment of genes exhibiting down-regulated expression in *fam168b* morphants at 96 hpf *via* KEGG pathway analysis (**C1**) and venn diagrams representing the overlapping down-regulated nervous system genes in *pou3f1*, *fam168b* and *fam168a* morphants at 96 hpf (**C2**). (**D**) Gene ontology (GO) classification of the genes exhibiting down-regulated expression in fam168b morphants at 96 hpf (**D1**) and venn diagrams representing the overlapping down-regulated synapse genes in *pou3f1*, *fam168b* and *fam168a* morphants at 96 hpf (**D2**). (**E**) WISH analysis of CNS myelin marker *mbp* in WT embryos, *fam168b*, *fam168a* and *pou3f1* morphants or mutants at 96 hpf (**E1-E6**), and quantification analysis of the WISH data in different samples (**E7-E8**). (**F**) qRT-PCR expression analysis of CNS myelin marker **mbp** in embryos injected with different morpholinos at 96 hpf. Each experiment was repeated three times, and a representative result is shown. Data are mean ± SD. **A, B, E,** lateral view, anterior to the left and dorsal to the up. The red arrow indicates mbp-expression in the spinal cord. \**P* < 0.05, \*\**P* < 0.01, \*\*\**P* < 0.001. NS, not significant. Scale bars, 200 μm (A, B) and 100 μm (E). See also Fig S4.

Loci in the *fam168b* promoter from -1672 to -1414, -1414 to -1240, and -1240 to -927 were obviously hypermethylated in Cu^2+^ stressed larvae (Fig 4A). Thus, we further investigated the roles of the aforementioned hypermethylated loci in regulating gene transcription. Different truncated promoter driven GFP fluoresces were almost distributed throughout the neural ectoderm in the injected embryos (Fig 4E), indicating their transcriptional activities during embryogenesis. Cu^2+^ slightly down-regulated the GFP fluoresce driven by different truncated promoters in embryos (Figs 4E2, E4, E6, E8 and S3D). Additionally, compared to GFP fluoresce driven by *fam168b* promoter from -1672 in the injected embryos (Figs 4E1, E2 and S3D), the GFP fluoresce from -1414 was obviously reduced (Figs 4E3, E4 and S3D), with a further reduction in the GFP fluoresce from -1240 (Figs 4E5, E6 and S3D) and -927 (Figs 4E7, E8 and S3D) truncated promoter driven GFP plasmids. The luciferase activity assays revealed that the gradient truncation of the *fam168b* promoter led to a gradual decrease of the transcriptional activity in the sequence of -927 promoter mutant < -1240 mutant <-1414 mutant < -1672 (Figs 4G and S3D). The schema of truncated *fam168b* promoter constructs was shown in Fig 4F.

**Fig 6.**
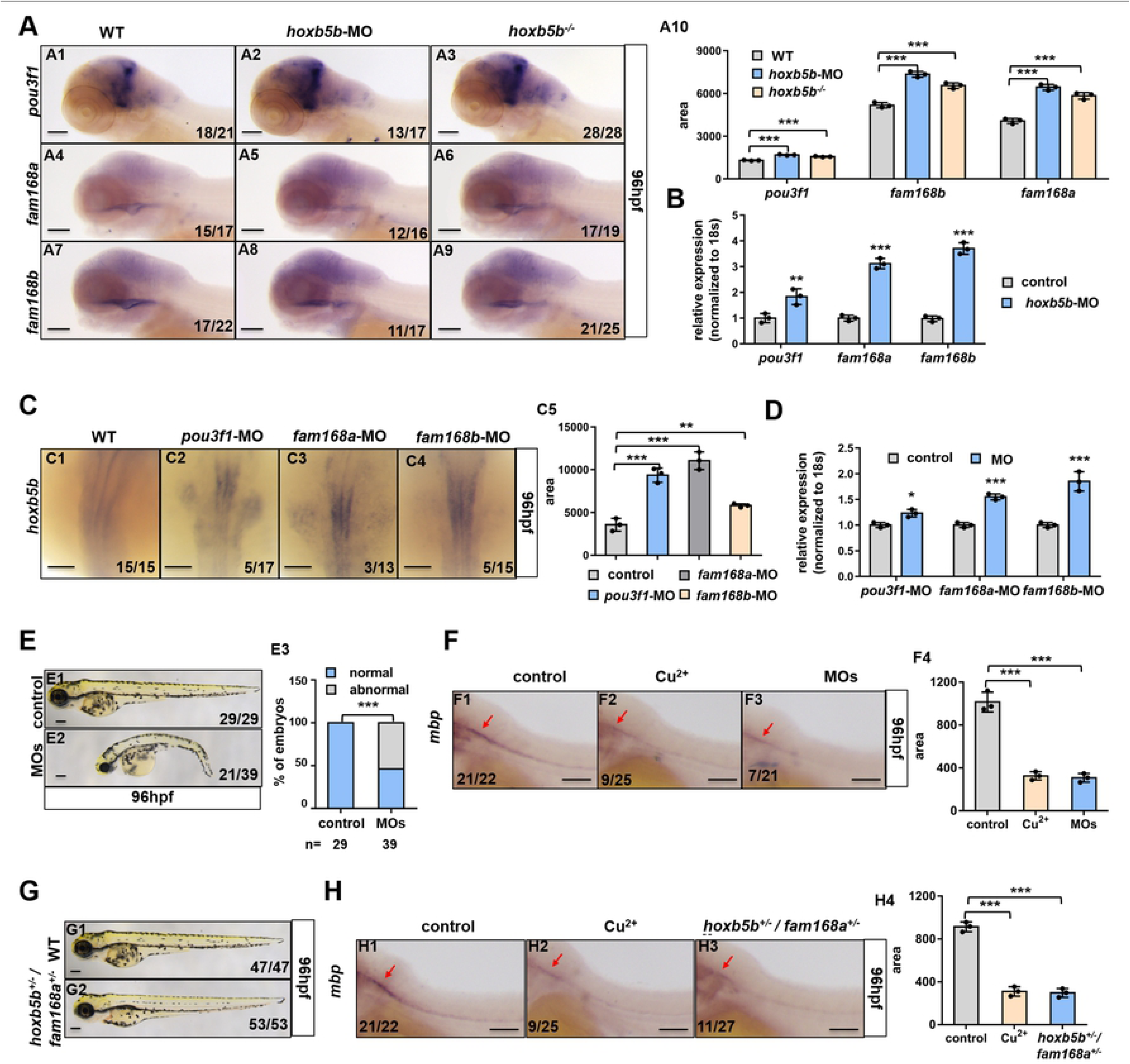
Wnt&Notch-*hoxb5b* signaling axis and *fam168a*/*fam168b*/*pou3f1* transcriptional factors in embryogenesis. (**A**)WISH analysis of hypermethylated genes *pou3f1*, *fam168a* and *fam168b* expression in the control or Cu^2+^ stressed larvae at 96 hpf in *hoxb5b*-MO injected- or *hoxb5b*^-/-^ embryos (**A1-A9**). Quantification analysis of the WISH data in different samples (**A10**). (**B**) qRT-PCR analysis of *pou3f1*, *fam168a* and *fam168b* expression in *hoxb5b*-MO injected embryos at 96 hpf. (**C**)WISH analysis of *hoxb5b* expression in *pou3f1*, *fam168a* or *fam168b* morphants at 24 hpf. Quantification analysis of the WISH data in different samples (**C5**). (**D**) qRT-PCR expression analysis of *hoxb5b* in *pou3f1*, *fam168a* or *fam168b* morphants at 24 hpf. (**E**) Phenotypes of embryos injected separately with *hoxb5b*, *pou3f1*, *fam168a* and *fam168b* MOs at 96 hpf, and the percentage of embryos exhibiting abnormal development in different samples (**E3**). (**F**) WISH analysis of CNS myelin marker *mbp* expression in the control, Cu^2+^ stressed or MOs injected embryos at 96 hpf. Quantification analysis of the WISH data in different samples (**F4**). (**G**) Phenotypes of *hoxb5b*^-/-^/*fam168a*^-/-^ embryos at 96 hpf. (**H**) WISH analysis of CNS myelin marker *mbp* expression in the control, Cu^2+^ stressed or hoxb5b^-/-^/fam168a^-/-^ embryos at 96hpf. Quantification analysis of the WISH data in different samples (**H4**). Each experiment was repeated three times, and a representative result is shown. Data are mean ± SD. **C**, dorsal view, anterior to the top, **A, E, F, G, H,** lateral view, anterior to the left and dorsal to the up. The red arrow indicates mbp-expression in the spinal cord. \**P* < 0.05, \*\**P* < 0.01, \*\*\**P* < 0.001. NS, not significant. Scale bars, 100 μm (A, C, F, H) and 100 μm (E, G). See also Fig S5.

### CNS myelin formation in *fam168a*/*fam168b* loss- and gain-of-function embryos

The function of *fam168a* and *fam168b* during embryogenesis was further tested by knockdown and knockout of *fam168a* and *fam168b* in embryos. The transcripts of *fam168a* and *fam168b* were distributed ubiquitously among the whole embryo at the early stages (Fig S4A and S4B). Their predominant expression in the brain was observed at 96 hpf (Figs 5A1 and A2), similar to the expression pattern of *pou3f1* in the embryos at this stage (Fig 5A3). The WT embryos injected with *fam168a* or *fam168b* MO exhibited similar developmental defects, such as shortened body, microcephalia, and slight ventralization at 24 hpf (Figs 5B2 and 5B3) and 96 hpf (Figs 5B6 and B7), similar to the developmental defects observed in *pou3f1* morphants (Fig 5B4 and B8). Meanwhile, *fam168a^-/-^* mutant with a 4-bp deletion (Fig S4C1) exhibited a normal-like phenotype (Fig 5B10) and *fam168b*^-/-^ mutant with 1-bp deletion (Fig S4C2) displayed microcephalia and slight ventralization at 96 hpf (Fig 5B11).

**Fig 7.**
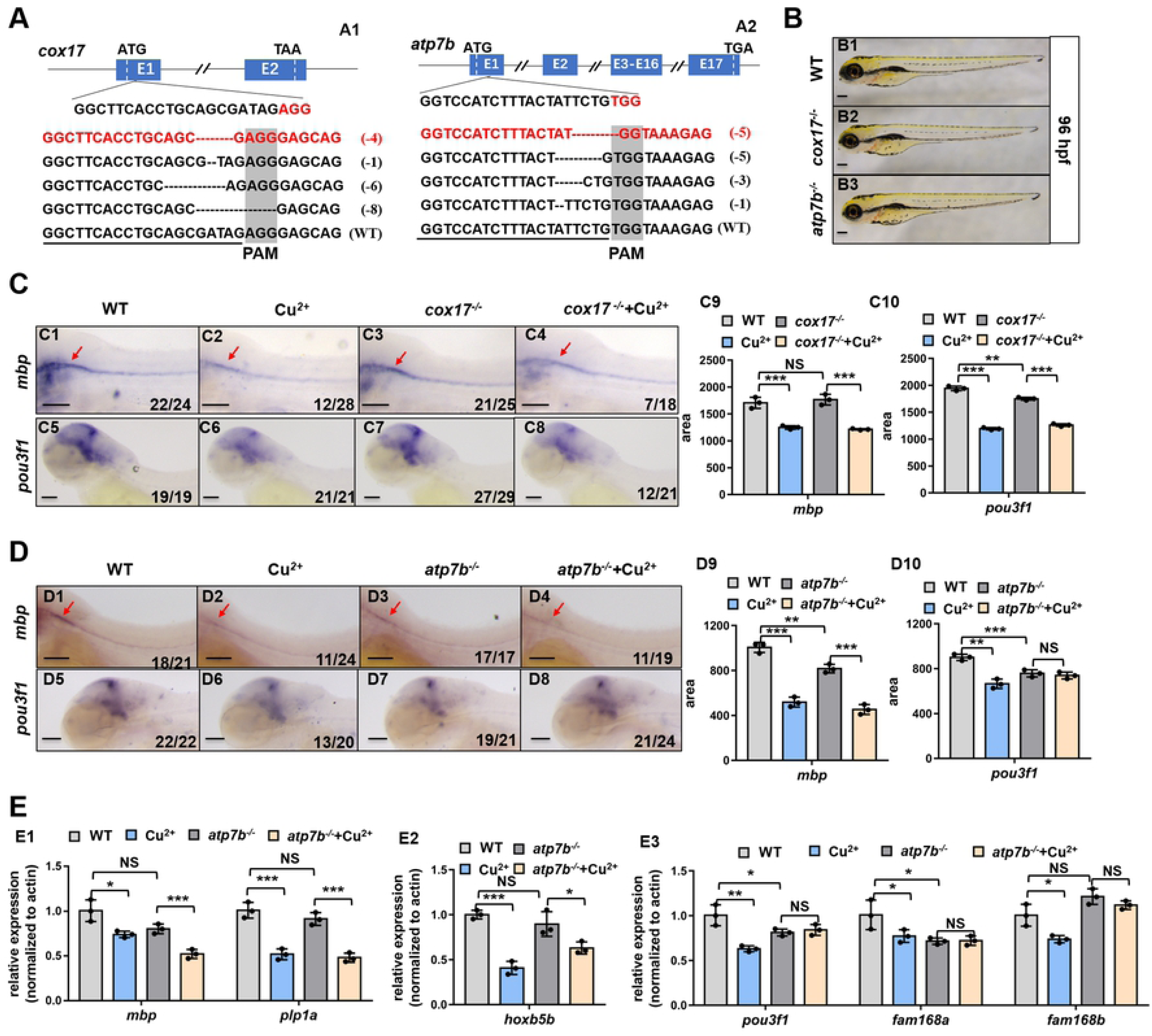
Myelin and axon formation in Cu^2+^ stressed *cox17* and *atp7b* mutant. (**A**) Schematic diagrams showing the genomic structure and genetic mutations of zebrafish genes *cox17* and *atp7b*, and the red line in gene *cox17* or *atp7b* respectively indicates the genotypic deletion of the mutation of *cox17* or *atp7a* used in this study. ATG denotes the translation start codon; TGA, the translation termination codon; PAM, the protospacer adjacent motif; slash, intron; blue horizontal bar, exon; dotted lines, the deletion of *cox17* and *atp7b*; numbers, the length of mutant base. (**B**) Phenotypes of *cox17* (**B2**) mutant with a 4-bp deletion and *atp7b* (**B3**) mutant with a 5-bp deletion at 96 hpf. (C) WISH analysis of the expression of CNS myelin marker *mbp* (**C1-C4**) and myelination transcriptional factor *pou3f1* (**C5-C8**) in WT embryos, Cu^2+^ stressed WT embryos, *cox17*^-/-^ mutant embryos or Cu^2+^ stressed *cox17*^-/-^ mutant at 96 hpf. Quantification analysis of the WISH data in different samples (**C9-C10**). (**D**) WISH analysis of the expression of CNS myelin marker *mbp* (**D1-D4**) and myelination transcriptional factor *pou3f1* (**D5-D8**) in WT embryos, Cu^2+^ stressed WT embryos, *atp7b*^-/-^ mutant embryos or Cu^2+^ stressed *atp7b*^-/-^ mutant at 96 hpf. Quantification analysis of the WISH data in different samples (**D9-D10**). (**E**) qRT-PCR expression analysis of the CNS myelin marker *mbp* (**E1**), transcriptional factor *hoxb5b* (**E2**), myelination transcriptional factor *pou3f1*, *fam168a* and *fam168b* (**E3**) in WT embryos, Cu^2+^ stressed WT embryos, *atp7b*^-/-^ mutants or Cu^2+^ stressed *atp7b*^-/-^ mutants at 96 hpf. Each experiment was repeated three times, and a representative result is shown. Data are mean ± SD. **B, C, D**, lateral view, anterior to the left and dorsal to the up. The red arrow indicates mbp-expression in the spinal cord. \**P* < 0.05, \*\**P* < 0.01, \*\*\**P* < 0.001. NS, not significant. Scale bars, 200 μm (A) and 100 μm (C, D). See also Fig S6.

**Fig 8.**
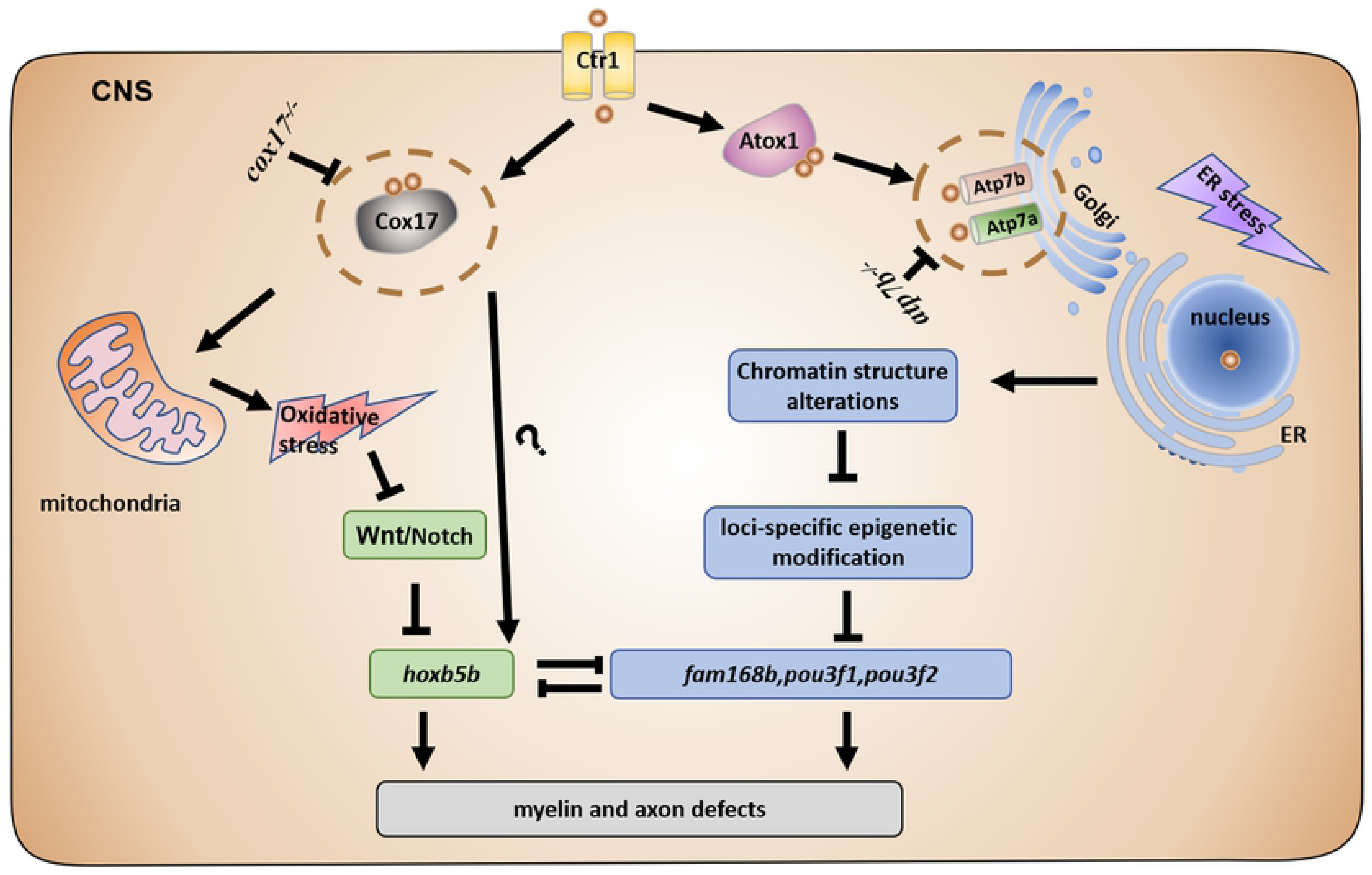
Working model of copper in inducing CNS defects. Cu^2+^ overloaded in the CNS cells of Cu^2+^ stressed WT, *cox17*, or *atp7b* null larvae, leading to the down-regulation of the Wnt/Notch-*hoxb5b* signaling axis in *atp7b* null larvae, the promoter methylation of genes *pou3f1*/*pou3f2*/*fam168b* and their down-regulated expression in *cox17* null mutants, and leading to the down-regulated expression of the two signaling pathways in WT embryos, then, resulted in CNS defects in copper stressed WT, *cox17*^-/-^, or *atp7b*^-/-^ larvae respectively.

Transcriptional profiles in *fam168a* and *fam168b* morphants were investigated by KEGG pathway (Figs 5C1, S4D1 and Table S10, S11) and cellular component GO analyses (Figs 5D1, S4E1 and Table S12, S13). They showed enrichment in the nervous system and synapse for the down-regulated DEGs, identical to transcriptional profiles in *pou3f1* morphants (Figs S4D2, S4E2, Table S14 and S15). Additionally, 85 genes in the nervous system (Fig 5C2) and 8 genes in synapse (Fig 5D2) were down-regulated and overlapped in the three *fam168a*, *fam168b*, and *pou3f1* morphants. Meanwhile, 104 genes in the nervous system (Fig S4D3) and 8 genes in synapse (Fig S4E3) were down-regulated and overlapped in both *fam168a* and *fam168b* morphants.

In this study, CNS myelin and axon development in *fam168a/b* loss/knockdown-of-function embryos were further tested in term of *mbp* and *plp1a*expression. *Mbp* and *plp1a* exhibited obviously reduced expression in both *fam168a/b* morphants and *fam168a* homozygous mutant by qRT-PCR and WISH assays (Figs 5E, 5F, S4F and S4G), similar to its expression in *pou3f1* morphants (Figs 5E and 5F) and in Cu*^2+^* stressed embryos.

Additionally, *fam168a, fam168b*, and *pou3f1* mRNA were injected separately into Cu^2+^ stressed embryos to test whether they could rescue the myelin formation. *Mbp* expression was partially recovered in the Cu*^2+^* stressed embryos injected separately with *fam168a, fam168b,* and *pou3f1* mRNA (Fig S4H).

### Wnt&Notch-*hoxb5b* signaling and *fam168a/fam168b/pou3f1* transcriptional factors in embryogenesis

The crosstalk between Wnt&Notch-*hoxb5b* and *fam168a*/*fam168b*/*pou3f1* transcriptional factors underlying Cu*^2+^*-induced myelin and axon developmental defects was explored separately by analysis of the expression of hypermethylated genes *pou3f1*, *fam168a*, and *fam168b* in *hoxb5b* morphants and *hoxb5b*^-/-^ mutants and vice versa. *Pou3f1*, *fam168a*, and *fam168b* showed significantly increased expression in both *hoxb5b* morphants and *hoxb5b*^-/-^ mutants at 96 hpf (Figs 6A, 6B and S5A). So did *hoxb5b* in *pou3f1*, *fam168a*, and *fam168b* morphants (Figs 6C, 6D and S5B).

Furthermore, we detected the combined effects of down-regulation of the two signaling pathways on the embryonic development and *mbp* expression. Morphants injected with the combined MOs of *hoxb5b*, *pou3f1*, *fam168a* and *fam168b* exhibited similar phenotypic defects (Fig 6E) and obviously reduced expression of CNS myelin marker *mbp* (Figs 6F and S5C). Meanwhile, *hoxb5b*^+/-^*fam168a*^+/-^ mutants exhibited normal-like phenotype at 96 hpf (Fig 6G), but an obviously reduced expression of CNS myelin marker *mbp* (Figs 6H and S5D).

### CNS myelin and axon formation in copper stressed *cox17^-/-^*, *atp7b^-/-^*, and *atp7a^-/-^* mutants

The question of in which organelle Cu*^2+^*overload resulted in the changed expression of the down-stream signaling and the CNS myelin and axon defects was investigated by using *cox17^-/-^* (Fig 7A1) and *atp7b^-/-^* (Fig 7A2) null mutants. *Cox17^-/-^* and *atp7b^-/-^* null mutants exhibited normal-like phenotypes at 96 hpf (Figs 7B). However, RNA-seq analysis revealed that genes in the nervous system (Fig S6A1), synapse (Fig S6A2), and axon (Fig S6A3) exhibited reduced expression in *cox17^-/-^* mutants.

Furthermore, the expression of the CNS myelin markers *mbp*, *plp1a* and genes *pou3f1*, and *fam168a*&*fam168b* was tested in *cox17^-/-^* or *atp7b^-/-^* mutants with and without Cu^2+^ stimulation. When compared with the WT control, *mbp* and *plp1a* showed no expression change in *cox17^-/-^* mutants, and so did *fam168a* and *fam168b* in *cox17^-/-^* mutants, but *pou3f1* exhibited a slightly reduced expression in *cox17^-/-^* mutants (Figs 7C and S6B). When compared with their expression in *cox17^-/-^* mutants without copper stimulation (Figs 7C3, C7, C9, C10 and Figs S6B3, B7, B11 B13 and B14), *mbp*, *plp1a*, *pou3f1*, *fam168a*, and *fam168b* exhibited reduced expression in Cu^2+^ stressed *cox17^-/-^* mutants at 96 hpf (Figs 7C4, C8, C9, C10 and Figs S6B4, B8, B12, B13 and B14), similar to their expression tendency in Cu^2+^ stressed WT embryos (Figs 7C1, C2, C5, C6, C9, C10 and Figs S6B1,B2,B5, B6, B9,B10, B13 and B14). The percentages of embryos with reduced expression of the aforementioned genes were significantly increased in either WT or *cox17^-/-^* mutants after Cu^2+^ stimulation (Fig S6C). Additionally, RT-PCR analysis also unveiled the significantly reduced expression of *mbp*, *plp1a* (Fig S6D1), *pou3f1*, *fam168a*, or *fam168b* (Fig S6D3) in either Cu^2+^ stressed WT or *cox17^-/-^* embryos, but no significant change of *hoxb5b* in Cu^2+^ stressed *cox17^-/-^* mutants (Fig S6D2).

Myelin specification was further tested in *atp7b^-/-^* embryos after copper stimulation. When compared with their expression in WT embryos, *mpb*, *plp1a*, *pou3f1*, *hoxb5b*, and *fam168a* exhibited significantly reduced expression in *atp7b^-/-^* embryos (Figs 7D, 7E, and Figs S6E-F), with the expression of *mbp* (Figs 7D4, D9, 7E1, and Fig S6F1), *plp1a* (Fig 7E1) and *hoxb5b* (Fig 7E2) being more significantly reduced in *atp7b*^-/-^ embryos after copper stimulation. However, WISH and qRT-PCR assays revealed no significant expression change in *pou3f1* and *fam168a*&*fam168b* in *atp7b*^-/-^ embryos after copper stimulation (Figs 7D8-D10, 7E3 and S6E, S6F2).

ER stress antagonist PBA was used to further study the role of copper-induced ER stresses in copper-induced down-regulated expression of *mbp*, *hoxb5b*, and *fam168a*. No significant recovery was observed in the expression of the three genes in Cu^2+^ stressed embryos after PBA co-treatment (Fig S6G).

## Discussion

Cu^2+^ has been reported to induce dysfunctional locomotor in zebrafish larvae(22), but the underlying mechanisms are still poorly understood. In this study, Cu^2+^ was revealed to induce uncompacted and thinner myelin in the spinal cord, which was consistent with the observations in *epb41l2* mutants with dysfunctional locomotor behaviors(36).

It is reported that *mbp*, a widely used marker for myelin(37), expressed in both the CNS and PNS myelin. *Olig2* expressed in oligodendrocyte and *olig2* driven fluoresce specifically labels oligodendrocytes and axon in the *olig2*:DsRed transgenic zebrafish line. In this study, *mbp* and *olig2* exhibited significantly reduced expression in the spinal cord, and the length of axon was significantly reduced in Cu^2+^ stressed embryos, indicating Cu^2+^ induced CNS myelin and axon defects in zebrafish. Additionally, the shortened axon might be the secondary damage of defective myelin formation in Cu^2+^ stressed larvae, as indicated by previous studies showing that myelin abnormalities might precede evidence of axonopathies(38, 39).

Cu^2+^ specifically induced the down-regulated expression of Wnt signaling and Notch signaling and their down-stream mediator *hoxb5b*, rather than other *hox* genes in zebrafish embryos(22). Inhibition of Wnt signaling has been shown to induce hypomyelination, whereas the activation of Wnt signaling significantly increased the transcription of myelin genes in mouse(29). Notch signaling has been revealed to regulate the differentiation of oligodendrocyte precursor cells, and influence oligodendrocyte maturation and myelin wrapping(30, 40), and *hox5* has been unveiled to regulate axon extension in motor neurons(26). Consistently, knockdown/out of *hoxb5b* zebrafish phenocopied the defective CNS myelin and axon observed in Cu^2+^ stressed embryos, whereas ectopic *hoxb5b* expression rescued the defects of CNS myelin and axon, indicating Cu^2+^ partially inhibited CNS myelin and axon marker expression *via* suppression of *hoxb5b*. The normal-like morphology of *hoxb5b^-/-^* mutant might result from the genetic compensation reported recently(41). Additionally, this study unveiled that both Wnt agonist BIO and Notch signaling activator NICD not only recovered the reduced expression of *hoxb5b*, but also recovered the myelin and axon defects in Cu^2+^ stressed embryos, further demonstrating that down-regulated expression of Wnt&Notch-*hoxb5b* signaling mediated Cu^2+^-induced myelin and axon developmental defects.

DNA methylation has been suggested to involve in regulation of gene expression and associate with a series of copper induced demyelinating diseases such as WD and AD (16, 17). In this study, it was unveiled that *pou3f1*, *pou3f2* and *fam168b* exhibited down-regulated expression but hypermethylation separately in their promoter in Cu^2+^ stressed embryos. Their promoter hypermethylation and reduced expression in Cu^2+^ stressed embryos suggested the potential correlation of gene transcription with its promoter methylation in Cu^2+^ stressed larvae.

In this study, it was shown that both *fam168b* and *fam168a* exhibited down-regulated expression in Cu^2+^ stressed embryos, with a highly similar expression pattern to that of *pou3f1* during fish embryogenesis. Additionally, similar transcriptional profiles and gene expression patterns, such as enrichment of nervous system and synapse for the down-regulated DEGs as well as down-regulated expression of CNS myelin genes, were observed in both *fam168a*&*fam168b* loss/knockdown-of-function embryos and *pou3f1* morphants. *Pou3f1* and *pou3f2* were critical transcription factors in the conversion of embryonic stem cells into neuron and glial cells(42), and the function of *pou3f2* was largely overlapped with that of *pou3f1* in driving the transition from promyelinating to myelinating cells(43). *Fam168b*, a novel neural gene identified recently, has been reported to control neuronal survival and differentiation as well as be specifically expressed in myelinated neuron in the CNS in human and mice(44, 45), to exhibit significantly down-regulated expression in AD brains (44), but has never been reported to be involved in myelin development. In this study, similar transcriptional profiles were observed in *fam168a*, *fam168b*, and *pou3f1* morphants, and ectopic expression of *fam168a*, *fam168b*, or *pou3f1* could rescue CNS myelin defects in Cu^2+^ stressed embryos. Taken together, *fam168a* and *fam168b* might be novel transcriptional factors similar to *pou3f1* in oligodendrocyte differentiation and the subsequent myelin cell development.

In this study, truncated promoter driven GFP and luciferase assays unveiled that the different hypermethylated loci in *fam168b* promoter, such as locus from -1672 to -1414, -1414 to -1240, and – 1240 to – 927, were critical for its transcriptional regulation during embryogenesis and in cells. The deletion of the aforementioned loci in *fam168b* promoter could induce significant down-regulation of its transcriptional activity, suggesting the hypermethylated loci are required and pivotal for *fam168b* transcriptional activation. Collectively, we demonstrated for the first time that Cu^2+^ might induce hypermethylation in the *fam168b* promoter, which is correlated with its down-regulated expression in Cu^2+^ stressed embryos. However, the down-regulated expression of *fam168b* occurred at 24 hpf, followed by hypermethylation of its promoter at 96 hpf in copper stressed embryos, suggesting the chromatin structure of transcriptional complex with its binding chromosome DNA might be damaged before promoter hypermethylation. This is consistent with the point in recent reports showing that regional methylation could be a secondary consequence of changes in transcriptional complex and chromosome DNA structure(34, 46).

Additionally, up-regulated expression of epigenetic mediators *pou3f1*/*fam168a*/*fam168b* was observed in *hoxb5b* loss/knockdown embryos, but significantly increased expression of *hoxb5b* was observed in *pou3f1*, *fam168a*, or *fam168b* morphants, not only suggesting *pou3f1*/*fam168a*/*fam168b* and *hoxb5b* acted in embryogenesis in a seesaw manner, but also indicating that hypermethylated *pou3f1* and *fam168a*&*b* were parallel factors to Wnt/Notch-*hoxb5b* signaling axis in mediating copper-induced myelin and axon defects.

The transfer of copper to mitochondria was assumed to be blocked in copper stressed *cox17^-/-^* mutant, and *cox17^-/-^* embryos fail to produce ROS after copper stimulation (47), but defects of CNS myelin and axon were still observed in this study, suggesting that copper-induced myelin and axon defects might not be essentially mediated by copper-induced ROS and by the function of *cox17* alone. Moreover, in this study, endoplasmic reticulum (ER) stress alleviant PBA was found unable to recover the expression of *mbp* in Cu^2+^ stressed WT embryos, suggesting copper-induced ER stresses might not alone mediate copper-induced CNS myelin development defects.

The *cox17^-/-^* larvae exhibited significantly reduced expression in *pou3f1*, *fam168a*, or *fam168b* after copper stimulation, while Cu^2+^ stressed *atp7b^-/-^* larvae exhibited slightly down-regulation in the expression of *pou3f1* but no expression change in *fam168a* and *fam168b*. However, *hoxb5b* exhibited significantly reduced expression in Cu^2+^ stressed *atp7b^-/-^* mutants but not in Cu^2+^ stressed *cox17^-/-^* mutants. This not only suggested that copper induced changes in the promoter chromatin structure and the down- regulated expression of the *fam168a*/*fam168b/pou3f* genes independent of the function of *cox17* alone, but also implying that copper required the integral function of *atp7b* rather than *cox17* to induce the promoter methylation and the resultant reduced expression of genes *pou3f1*/*fam168a*/*fam168b*, and required integral function of *cox17* rather than *atp7b* for the down-regulated expression of Wnt&Notch- *hoxb5b* axis. This might help to explain why myelin defects still occurred in either Cu^2+^ stressed *cox17^-/-^* or *atp7b^-/-^* embryos. In this study, we demonstrated that the epigenetic methylation of *pou3f1*/*fam168a*/*fam168b* in Cu^2+^ stressed *cox17^-/-^* embryos or the down-regulated expression in the Wnt&Notch-*hoxb5b* axis in Cu^2+^ stressed *atp7b^-/-^* embryos separately mediated the down-regulated expression of myelin genes in the Cu^2+^ stressed mutants. It has been unveiled that copper could locate in cell nucleus and damage the chromatin structure directly (48, 49). Thus, this study provided the direct evidence for the first time that copper damages chromatin structure independent of ROS in DNA methylation during fish embryogenesis.

In summary, this study confirmed the structural and detailed molecular characters of CNS myelin and axon defects occurring in copper stressed embryos. It was shown that copper induced ROS and led to down-regulation of Wnt&Notch-*hoxb5b* axis, with copper directly inducing locus-specific methylation and the down-regulated expression of *pou3f1*/*fam168a*/*fam168b* genes to mediate myelin and axon defects in copper stressed embryos. The working model is illustrated in Fig 8 for an intuitive understanding of how copper induces CNS defects. The combined data from the current study added novel insights into the mechanisms underlying the unbalanced copper homeostasis in cells linking with neurological disorders.

## Materials and methods

### Ethics statement

All experiments involving fish in this study were performed in accordance with the recommendations in the Guide for the care and use of Laboratory Animals of the Ministry of Science and Technology of China, which was approved by the Scientific Committee of Huazhong Agricultural University (permit number HZAUFI-2016-007).

### Fish stocks

Wild-type zebrafish (*Danio rerio*) (AB) maintenance, breeding and staging were performed as described previously(50). *Tg*(*olig2*:dsRED) and *Tg*(*mbp*:EGFP) transgenic lines were obtained from China Zebrafish Resource Center (http://www.zfish.cn/), and the catalog numbers of the lines used were listed in Table S1.

### Morpholinos and Cas9/gRNA

The CRISPR/Cas9 genome editing system was reported as an effective tool for gene editing in organisms (51, 52). In this study, the CRISPR/Cas9 system was used to construct *homeobox B5b* (*hoxb5b*), *family with sequence similarity 168 member A* (*fam168a*), *family with sequence similarity 168 member B* (*fam168b*), *ATPase copper transporting alpha* (*atp7a*), *ATPase copper transporting beta* (*atp7b*) and *cytochrome c oxidase copper chaperone COX17* (*cox17*) mutants. The guide RNAs (gRNAs) were designed to target the first exon of aforementioned genes by ZiFiT Targeter Version 4.2 at the following URL (http://zifit.partners.org/ZiFiT/CSquare9Nuclease.aspx). Sequences of gRNAs are listed in Table S2. The morpholinos (MOs), including *hoxb5b*-MO, *pou3f1*-MO, *fam168a*-MO, and *fam168b*- MO, were purchased from Gene Tools, LLC (Philomath, Oregon, USA) and their sequences are listed in Table S3.

### Drug exposure

Copper and 6-Bromoindirubin-3′-oxime (BIO) (Sigma-Aldrich, USA) were prepared as described previously (22, 27). Zebrafish embryos developed to sphere stage (4 hpf, hours post fertilization) or early were exposed to 3.9 μM copper at random. BIO was added at bud stage. Embryos were harvested at indicated stages. Each group was biologically repeated 3 times.

### Transmission electron microscope (TEM) analysis

TEM was performed to test CNS myelin structure in the control and copper stressed embryos at 5 dpf (days post fertilization). A transmission electron microscope (Hitachi H-7650 TEM Japan) was used to acquire the images. G-ratio (axon diameter/myelinated fiber diameter) was calculated to assess myelin thickness. A lower g-ratio indicated a greater myelin thickness. The axon diameter and myelinated fiber diameter were measured using the image J software (NIH, Bethesda, Maryland).

### Plasmid construction

The full-length *hoxb5b*, *fam168a*, *fam168b*, *POU class 3 homeobox 1* (*pou3f1*), and the intracellular domain of *notch receptor 3* (*notch3*) (NICD) were amplified using the primers shown in Table S4 and cloned into pCS2 vector for synthesizing mRNAs. 5’ unidirectional deleted mutants of *fam168b* promoter, including -1672, -1414, -1240, -927, -623, and -284, were amplified using the primers shown in Table S5 and cloned separately into pCS2-GFP vector and pGL3 vector. All constructs were verified by sequencing.

### mRNA Synthesis and Injection

For mRNA preparation, capped mRNAs were synthesized using the mMessage mMachine kit (Ambion) according to the manufacturer’s instructions. The synthesized mRNAs were diluted into different concentrations and injected into one-cell stage embryos as reported previously(53).

### Quantitative RT-PCR analysis

Zebrafish embryos were collected at indicated stages. Total RNA was isolated from 50 whole embryos/sample using Trizol reagent (Invitrogen). cDNA was synthesized using a M-MLV Reverse-Transcript Kit (Applied Biological Materials Inc, BC, Canada). qRT-PCR was performed as described previously(22, 53, 54). The sequences of the RT-PCR primers were listed in Table S6.

### Whole-mount in situ hybridization

Probes for zebrafish *myelin basic protein a* (*mbp*), *oligodendrocyte lineage transcription factor 2* (*olig2*), *hoxb5b*, *pou3f1*, *fam168a*, and *fam168b* were amplified from cDNA pools using primers shown in Table S7. Whole-mount in situ hybridization (WISH) was performed as described previously(50, 53, 55). WISH embryos were photographed with a Leica M205FA stereomicroscope. The signal area in each image was calculated by Image J software (NIH, Bethesda, Maryland). Embryos with changed expressions in the tested genes were identified and their percentage was calculated as reported in our previous works(22, 53, 56).

### RNA-sequencing (RNA-Seq) and analysis

WT embryos, WT embryos injected with a different MO of *hoxb5b*, *pou3f1*, *fam168a*, or *fam168b*, *cox17^-/-^* and *atp7a^-/-^* mutants and Cu^2+^ stressed *atp7a^-/-^* mutants at 96 hpf were lysed by Trizol reagent (Ambion, Life Technologies) for RNA preparation. The RNAs were then reversely transcribed, and amplified cDNA were sequenced on a BGISEQ-500 platform (BGI, Wuhan, China). Quantile normalization and subsequent data processing were performed using the RSEM v1.2.8 software package. Pathway and GO (Gene ontology) analyses were carried out to determine the roles of the differentially expressed genes (DEGs). The Hierarchical Cluster Tree (dendrogram) was constructed to show the relationships of expression levels among different samples.

### Confocal microscopy

Embryos were anesthetized with a low dose of tricaine and mounted on dishes with 1% low-melting agarose. Confocal images were captured by a Leica (Wetzlar, Germany) TCS SP8 confocal laser microscope. The fluorescence intensity of the positive cells in embryos was analyzed by software of Image J. Axon tracing and measurement was performed by using the Neuron J (Image J) software (National Institutes of Health, Bethesda, MD).

### Bisulfite PCR validation

Whole genome bisulfite sequencing (WGBS) has been performed in the control and the Cu^2+^ stressed embryos at 96 hpf, and 57 differential methylated genes (DMGs) were unveiled (35). In this study, the regions for differentially methylated loci of the candidate genes such as *fam168b*between the control group and Cu^2+^ stressed group were used for bisulfite PCR to validate the results of whole-genome bisulfite sequencing. The target fragments were amplified using specific primers (Table S8) designed with Methyl Primer Express v1.0 (http://www.urogene.org/cgi-bin/methprimer/methprimer.cgi). The obtained PCR products were purified using Min Elute Gel Extraction kit (OMEGA) and cloned into the pMD19-T Vector (Takara). The positive clones were confirmed by PCR and 12 clones were sequenced for each subject.

### Luciferase reporter assay

Different truncated mutant promoters of *fam168b* were used for luciferase assays in this study. The luciferase reporter assays were performed as described previously (53). The data were reported as the mean ± SD of three independent experiments in triplicate (53).

### Statistical analyses

The sample size used for different experiments in each group was larger than 10 embryos (n>10), and 2-3 biological replicates were performed for each test. Percentage analysis of the results among different groups was performed using hypergeometric distribution in the R-console software(57). Statistical data of the signal area and fluorescence level in different samples were analyzed using t-test by GraphPad Prism 7.00 software. Each dot represents signal level in an individual embryo. Statistical data of axon length were processed by GraphPad Prism 7.00 software. Each dot represents the length of each axon. The qRT-PCR data were analyzed by one-way analysis of variance (ANOVA) and post hoc Tukey’s test in the Statistic Package for Social Science (SPSS) 19.0 software. Each dot represents one repeat. The statistical analysis for luciferase reporter assay results was performed using GraphPad Prism 7.00 software (unpaired t-test) (GraphPad Software Inc). Data were presented as mean ± SD, \**P* < 0.05, \*\**P* < 0.01, \*\*\**P* < 0.001.

## Supplementary Information

Supplementary materials include 6 supplementary figures and figure legends, and 15 supplementary tables.

## Acknowledgments

We are grateful to Dr. Yibing Zhang (Institute of Hydrobiology, Chinese Academy of Science) for the gift of HEK293T cells. We thank Mr. Qinhan Xu, Bei Cui and Haojie Sun (Huazhong Agricultural University) for participating in the study.

## Author contributions

J.X.L conceived the project and wrote the manuscript; T.Z performed most of the experiments, analyzed data and wrote the manuscript. P.P.G and G.Z contributed to analyzed data and experiments. G.L.L, Y.P.F, H.F and J.F.G supervised the project and approved the final manuscript.

## Supporting Information Legends

**Fig S1. Expression of central neural system (CNS) myelin genes in Cu2+ stressed embryos. Related to Fig 1.**

**(A)**WISH analysis of expression for myelin oligodendrocyte marker *olig2* at 48hpf **(A1-A2)** in the control or Cu^2+^ stressed embryos, and the quantification analysis of the WISH data in different samples **(A3)**. **(B)** The percentage of embryos exhibited reduced expression of indicated CNS myelin genes in different samples. Each experiment was repeated three times, and a representative result is shown. Data are mean ± SD. **A1-A2**, lateral view, anterior to the left and dorsal to the up. The red arrow indicates *olig2*-expressing in oligodendrocyte. \**P* < 0.05, \*\**P* < 0.01, \*\*\**P* < 0.001. NS, not significant. Scale bars, 100 μm.

**Fig S2 Cu^2+^ induces CNS myelin and axon defects by down-regulating Wnt & Notch - *hoxb5b* regulatory axis in zebrafish embryos. Related to Fig 2 and 3.**

**(A)** The percentage of embryos exhibited reduced expression of indicated CNS myelin genes in WT embryos, *hoxb5b* Morphants, or *hoxb5b^-/-^* mutants at 96hpf**(A1-A2)**. **(A3)** The percentage of embryos exhibited reduced expression of *mbp* in the control, Cu^2+^ stressed embryos, or Cu^2+^ stressed embryos with ectopic *hoxb5b* expression at 96hpf. **(B)** Clustering analysis of Notch signaling genes with reduced expression in Cu^2+^ stressed embryos at 24 hpf**(B1)**. qRT-PCR analysis of Notch signaling genes *notch1a*, *dlc* and *mib* expression in the control or Cu^2+^ stressed embryos at 24gpf **(B2)**. The percentage of embryos exhibiting reduced expression of mbp in the control, Cu^2+^ exposed, or Cu^2+^ and BIO co-exposed embryos with NICD ectopic expression at 96 hpf**(B3)**. Each experiment was repeated three times, and a representative result is shown. Data are mean ± SD. \**P* < 0.05, \*\**P* < 0.01, \*\*\**P* < 0.001. NS, not significant.

**Fig S3 DNA methylation and transcriptional activity of myelin genes in Cu^2+^ stressed embryos. Related to** **Fig 4**.

**(A)** Analysis of bisulfite sequencing data for *fam168b*, *pou3f1* and *pou3f2* methylation levels in the control or Cu^2+^ stressed embryos**(A1)**. RNA-seq analysis of *fam168a*, *pou3f1* and *pou3f2b* expression in the control or Cu^2+^ stressed embryos**(A2)**.The percentage of embryos exhibited reduced expression of *pou3f1* in the control or Cu^2+^ stressed embryos at 96hpf**(A3)**.**(B)** Graphical representation of methylation patterns in the promoter domain of genes *pou3f1* **(B1)** and *pou3f2* **(B2)** separately in the control or Cu^2+^ stressed embryos. **(C)** Bisulfite PCR validation of *fam168b* in the control or Cu^2+^ stressed embryos. **(D)** Quantification of fluorescence level in 24hpf WT embryos injected separately with plasmid containing different truncated *fam168b* promoter driven GFP reporters. Each experiment was repeated three times, and a representative result is shown. Data are mean ± SD. *P < 0.05, **P < 0.01, ***P < 0.001. NS, not significant.

**Fig S4 Myelin and axon formation in *fam168a*/*fam168b* loss- and gain-of-function embryos Related to** **Fig 5**.

**(A)** The expression pattern of zebrafish *fam168a* during embryogenesis. **(B)** The expression pattern of zebrafish *fam168b* during embryogenesis. **(C)** Schematic diagram showing the genomic structure and a genetic mutation of *fam168a* **(C1)** and *fam168b* **(C2)** used in this study. ATG denotes the translation start codon; TGA denotes the translation terminate codon; PAM denotes the protospacer adjacent motif; slash denotes intron; blue horizontal bar denotes exon; dotted lines denote deletion of *fam168a* and *fam168b*; Numbers denote the length of mutant base. **(D)** Enrichment of genes exhibited down-regulated expression in *fam168a* or *pou3f1* morphants at 96hpf *via* KEGG pathway analysis **(D1-D2)** and venn diagrams representing the overlapping down-regulated nervous system genes in *fam168a* and *fam168b* morphants at 96 hpf **(D3)**. **(E)** Gene ontology (GO) classification of the genes exhibited down-regulated expression in *fam168a***(E2)** or *pou3f1* morphants at 96hpf **(E2)** and venn diagrams representing the overlapping down-regulated synapse genes in *fam168a* and *fam168b* morphants at 96 hpf **(E3)**. **(F)**WISH analysis of expression for CNS myelin marker *plp1a* at 96hpf in the WT, *fam168a*^-/-^ or *fam168b*^+/-^ embryos **(F1-F3)**, and the quantification analysis of the WISH data in different samples **(F4)**. **(G)** The percentage of embryos exhibiting reduced expression in different samples. **(H)** WISH analysis of CNS myelin marker *mbp* expression in the control, Cu^2+^ stressed embryos, or Cu^2+^ stressed embryos with ectopic different mRNA expression**(H1-H5)**. **(H6)** Quantification analysis of the WISH data in different samples. **(H7)** Percentage of embryos exhibited reduced expression of *mbp* in different samples. Each experiment was repeated three times, and a representative result is shown. Data are mean ± SD. **A1-A6, B1-B6**, dorsal view, anterior to the top, **A7-A8**, **B7-B8**, **F1-3, H** lateral view, anterior to the left and dorsal to the up. *P < 0.05, **P < 0.01, ***P < 0.001. NS, not significant. Scale bars, 100 μm.

**Fig S5 percentage of embryos exhibited abnormal expression in different samples. Related to Fig 6.**

**Fig S6 RNA-seq,WISH and qRT-PCR analysis unveils differentially expressed nervous system genes in *cox17* and *atp7b* mutant embryos. Related to** **Fig 7**.

**(A)** Clustering analysis of endoplasmic reticulum genes exhibited down-regulated expression in *atp7a^-/-^* mutant embryos at 96 hpf**(A1)**. Clustering analysis of the expression of endoplasmic reticulum stress genes in WT embryos, Cu^2+^ stressed WT embryos, *atp7a*^-/-^ mutants, or Cu^2+^ stressed *atp7a*^-/-^ mutants at 96hpf **(A2)**. Clustering analysis of myelin genes in WT embryos, Cu^2+^ stressed WT embryos, *atp7a^-/-^* mutants, or Cu^2+^ stressed *atp7a^-/-^* mutants at 96hpf **(A3). (B)** WISH analysis of the myelin genes *plp1a*, *fam168a* and *fam168b* expression in WT embryos, Cu^2+^ stressed WT embryos, *cox17^-/-^* mutant embryos or Cu^2+^ stressed *cox17^-/-^* mutant at 96hpf **(B1-B12)**. Quantification analysis of the WISH data in different samples **(B13-B14). (C)** Percentage of embryos exhibited reduced expression in different samples. **(D)** qRT-PCR analysis of CNS myelin markers *mbp*, *plp1a***(D1)**, transcriptional factor *hoxb5b***(D2)**, *pou3f*, *fam168a* and *fam168b***(D3)** expression in WT embryos, Cu^2+^ stressed WT embryos, *cox17^-/-^* mutant embryos or Cu^2+^ stressed *cox17^-/-^* mutant at 96hpf. **(E)** WISH analysis of myelination transcriptional factors *fam168a* and *fam168b* in WT embryos, Cu^2+^ stressed WT embryos, *atp7b^-/-^* mutant embryos or Cu^2+^ stressed *atp7b^-/-^* mutant at 96hpf **(E1-E8)**. Quantification analysis of the WISH data in different samples **(E9). (F)** Percentage of embryos exhibited reduced expression in different samples. **(H)** qRT-PCR analysis of *mbp* **(H1)**, *hoxb5b* **(H1)** and *fam168a* **(H3)** in WT embryos, Cu^2+^ stressed, and Cu^2+^ and PBA co-exposed embryos at 96hpf. Each experiment was repeated three times, and a representative result is shown. Data are mean ± SD. **B1-A16**, **F1-F8,** lateral view, anterior to the left and dorsal to the up. The red arrow indicates *mbp*-expressing in the spinal cord. *P < 0.05, **P < 0.01, ***P < 0.001. NS, not significant. Scale bars, 100 μm.

## References

1. Villegas R, Martin SM, O’Donnell KC, Carrillo SA, Sagasti A, Allende ML. Dynamics of degeneration and regeneration in developing zebrafish peripheral axons reveals a requirement for extrinsic cell types. Neural Dev. 2012;7:19.

2. Brewer GJ. Copper toxicity in the general population. Clin Neurophysiol. 2010;121(4):459–60.

3. De Boeck G, van der Ven K, Hattink J, Blust R. Swimming performance and energy metabolism of rainbow trout, common carp and gibel carp respond differently to sublethal copper exposure. Aquat Toxicol. 2006;80(1):92–100.

4. Sandahl JF, Baldwin DH, Jenkins JJ, Scholz NL. Odor-evoked field potentials as indicators of sublethal neurotoxicity in juvenile coho salmon (Oncorhynchus kisutch) exposed to copper, chlorpyrifos, or esfenvalerate. Can J Fish Aquat Sci. 2004;61(3):404–13.

5. Cheng MY, Ho HH, Huang TK, Chuang CF, Chen HY, Chung HW, et al. A compartmentalized culture device for studying the axons of CNS neurons. Anal Biochem. 2017;539:11–21.

6. Conforti L, Gilley J, Coleman MP. Wallerian degeneration: an emerging axon death pathway linking injury and disease. Nat Rev Neurosci. 2014;15(6):394–409.

7. Bauer NG, Richter-Landsberg C, Ffrench-Constant C. Role of the Oligodendroglial Cytoskeleton in Differentiation and Myelination. Glia. 2009;57(16):1691–705.

8. Simons M, Nave KA. Oligodendrocytes: Myelination and Axonal Support. Cold Spring Harb Perspect Biol. 2015;8(1):a020479.

9. Saab AS, Tzvetanova ID, Nave KA. The role of myelin and oligodendrocytes in axonal energy metabolism. Curr Opin Neurobiol. 2013;23(6):1065–72.

10. Gold BT, Johnson NF, Powell DK, Smith CD. White matter integrity and vulnerability to Alzheimer’s disease: preliminary findings and future directions. Biochim Biophys Acta. 2012;1822(3):416–22.

11. Takahashi N, Sakurai T, Davis KL, Buxbaum JD. Linking oligodendrocyte and myelin dysfunction to neurocircuitry abnormalities in schizophrenia. Prog Neurobiol. 2011;93(1):13–U7.

12. Liu J, Dietz K, DeLoyht JM, Pedre X, Kelkar D, Kaur J, et al. Impaired adult myelination in the prefrontal cortex of socially isolated mice. Nat Neurosci. 2012;15(12):1621–3.

13. Poggi G, Boretius S, Mobius W, Moschny N, Baudewig J, Ruhwedel T, et al. Cortical Network Dysfunction Caused by a Subtle Defect of Myelination. Glia. 2016;64(11):2025–40.

14. Saab AS, Nave KA. Myelin dynamics: protecting and shaping neuronal functions. Curr Opin Neurobiol. 2017;47:104–12.

15. Colman DR, Kreibich G, Frey AB, Sabatini DD. Synthesis and Incorporation of Myelin Polypeptides into Cns Myelin. J Cell Biol. 1982;95(2):598–608.

16. Ryu HW, Lee DH, Won HR, Kim KH, Seong YJ, Kwon SH. Influence of toxicologically relevant metals on human epigenetic regulation. Toxicol Res. 2015;31(1):1–9.

17. Medici V, Shibata NM, Kharbanda KK, LaSalle JM, Woods R, Liu S, et al. Wilson’s disease: changes in methionine metabolism and inflammation affect global DNA methylation in early liver disease. Hepatology. 2013;57(2):555–65.

18. Mordaunt CE, Shibata NM, Kieffer DA, Czlonkowska A, Litwin T, Weiss KH, et al. Epigenetic changes of the thioredoxin system in the tx-j mouse model and in patients with Wilson disease. Hum Mol Genet. 2018;27(22):3854–69.

19. Mastronardi FG, Noor A, Wood DD, Paton T, Moscarello MA. Peptidyl argininedeiminase 2 CpG island in multiple sclerosis white matter is hypomethylated. J Neurosci Res. 2007;85(9):2006–16.

20. Emery B, Lu QR. Transcriptional and Epigenetic Regulation of Oligodendrocyte Development and Myelination in the Central Nervous System. Cold Spring Harb Perspect Biol. 2015;7(9):a020461.

21. Dorts J, Falisse E, Schoofs E, Flamion E, Kestemont P, Silvestre F. DNA methyltransferases and stress-related genes expression in zebrafish larvae after exposure to heat and copper during reprogramming of DNA methylation. Sci Rep. 2016;6:34254.

22. Zhang T, Xu L, Wu JJ, Wang WM, Mei J, Ma XF, et al. Transcriptional Responses and Mechanisms of Copper-Induced Dysfunctional Locomotor Behavior in Zebrafish Embryos. Toxicol Sci. 2015;148(1):299–310.

23. Vonk WIM, Wijmenga C, van de Sluis B. Relevance of animal models for understanding mammalian copper homeostasis. Am J Clin Nutr. 2008;88(3):840s–5s.

24. Kaler SG. ATP7A-related copper transport diseases-emerging concepts and future trends. Nat Rev Neurol. 2011;7(1):15–29.

25. Schmidt K, Ralle M, Schaffer T, Jayakanthan S, Bari B, Muchenditsi A, et al. ATP7A and ATP7B copper transporters have distinct functions in the regulation of neuronal dopamine--hydroxylase. J Biol Chem. 2018;293(52):20085–98.

26. Philippidou P, Walsh CM, Aubin J, Jeannotte L, Dasen JS. Sustained Hox5 gene activity is required for respiratory motor neuron development. Nat Neurosci. 2012;15(12):1636–44.

27. Xu JP, Zhang RT, Zhang T, Zhao G, Huang Y, Wang HL, et al. Copper impairs zebrafish swimbladder development by down-regulating Wnt signaling. Aquat Toxicol. 2017;192:155–64.

28. Park HC, Appel B. Delta-Notch signaling regulates oligodendrocyte specification. Development. 2003;130(16):3747–55.

29. Tawk M, Makoukji J, Belle M, Fonte C, Trousson A, Hawkins T, et al. Wnt/beta-Catenin Signaling Is an Essential and Direct Driver of Myelin Gene Expression and Myelinogenesis. J Neurosci. 2011;31(10):3729–42.

30. Titus HE, Lopez-Juarez A, Silbak SH, Rizvi TA, Bogard M, Ratner N. Oligodendrocyte RasG12V expressed in its endogenous locus disrupts myelin structure through increased MAPK, nitric oxide, and notch signaling. Glia. 2017;65(12):1990–2002.

31. Hortopan GA, Baraban SC. Aberrant Expression of Genes Necessary for Neuronal Development and Notch Signaling in an Epileptic mind bomb Zebrafish. Dev Dynam. 2011;240(8):1964–76.

32. Lengerke C, Schmitt S, Bowman TV, Jang IH, Maouche-Chretien L, McKinney-Freeman S, et al. BMP and Wnt specify hematopoietic fate by activation of the Cdx-Hox pathway. Cell Stem Cell. 2008;2(1):72–82.

33. Medici V, Kieffer DA, Shibata NM, Chima H, Kim K, Canovas A, et al. Wilson Disease: Epigenetic effects of choline supplementation on phenotype and clinical course in a mouse model . Epigenetics-Us. 2016;11(11):804–18.

34. Radford EJ, Ito M, Shi H, Corish JA, Yamazawa K, Isganaitis E, et al. In utero effects. In utero undernourishment perturbs the adult sperm methylome and intergenerational metabolism. Science. 2014;345(6198):1255903.

35. Tai Z, Guan P, Wang Z, Li L, Zhang T, Li G, et al. Common responses of fish embryos to metals: an integrated analysis of transcriptomes and methylomes in zebrafish embryos under the stress of copper ions or silver nanoparticles. Metallomics. 2019;11(9):1452–64.

36. Saitoh Y, Ohno N, Yamauchi J, Sakamoto T, Terada N. Deficiency of a membrane skeletal protein, 4.1G, results in myelin abnormalities in the peripheral nervous system. Histochem Cell Biol. 2017;148(6):597–606.

37. Monk KR, Talbot WS. Genetic dissection of myelinated axons in zebrafish. Curr Opin Neurobiol. 2009;19(5):486–90.

38. Bartzokis G. Alzheimer’s disease as homeostatic responses to age-related myelin breakdown. Neurobiol Aging. 2011;32(8):1341–71.

39. Desai MK, Sudol KL, Janelsins MC, Mastrangelo MA, Frazer ME, Bowers WJ. Triple-transgenic Alzheimer’s disease mice exhibit region-specific abnormalities in brain myelination patterns prior to appearance of amyloid and tau pathology. Glia. 2009;57(1):54–65.

40. Rabadan MA, Cayuso J, Le Dreau G, Cruz C, Barzi M, Pons S, et al. Jagged2 controls the generation of motor neuron and oligodendrocyte progenitors in the ventral spinal cord. Cell Death Differ. 2012;19(2):209–19.

41. Ma ZP, Zhu PP, Shi H, Guo LW, Zhang QH, Chen YN, et al. PTC-bearing mRNA elicits a genetic compensation response via Upf3a and COMPASS components. Nature. 2019;568(7751):259–+.

42. Zhu Q, Song L, Peng G, Sun N, Chen J, Zhang T, et al. The transcription factor Pou3f1 promotes neural fate commitment via activation of neural lineage genes and inhibition of external signaling pathways. Elife. 2014;3.

43. Lin YMJ, Hsin IL, Sun HS, Lin S, Lai YL, Chen HY, et al. NTF3 Is a Novel Target Gene of the Transcription Factor POU3F2 and Is Required for Neuronal Differentiation. Mol Neurobiol. 2018;55(11):8403–13.

44. Mishra M, Akatsu H, Heese K. The novel protein MANI modulates neurogenesis and neurite-cone growth. J Cell Mol Med. 2011;15(8):1713–25.

45. Mishra M, Lee S, Lin MK, Yamashita T, Heese K. Characterizing the neurite outgrowth inhibitory effect of Mani. Febs Lett. 2012;586(19):3018–23.

46. Stadler MB, Murr R, Burger L, Ivanek R, Lienert F, Scholer A, et al. DNA-binding factors shape the mouse methylome at distal regulatory regions (vol 480, pg 490, 2011). Nature. 2012;484(7395):550-.

47. Zhang YJ, Zhang RT, Sun HJ, Chen Q, Yu XD, Zhang T, et al. Copper inhibits hatching of fish embryos via inducing reactive oxygen species and down-regulating Wnt signaling. Aquat Toxicol. 2018;205:156–64.

48. Cao H, Wang Y. Quantification of oxidative single-base and intrastrand cross-link lesions in unmethylated and CpG-methylated DNA induced by Fenton-type reagents. Nucleic Acids Res. 2007;35(14):4833–44.

49. Goswami S, Sanyal S, Chakraborty P, Das C, Sarkar M. Interaction of a common painkiller piroxicam and copper-piroxicam with chromatin causes structural alterations accompanied by modulation at the epigenomic/genomic level. Bba-Gen Subjects. 2017;1861(8):2048–59.

50. Liu JX, Hu B, Wang Y, Gui JF, Xiao WH. Zebrafish eaf1 and eaf2/u19 Mediate Effective Convergence and Extension Movements through the Maintenance of wnt11 and wnt5 Expression. J Biol Chem. 2009;284(24):16679–92.

51. Rosenbluh J, Xu H, Harrington W, Gill S, Wang X, Vazquez F, et al. Complementary information derived from CRISPR Cas9 mediated gene deletion and suppression. Nat Commun. 2017;8:15403.

52. Varshney GK, Carrington B, Pei WH, Bishop K, Chen ZL, Fan CX, et al. A high-throughput functional genomics workflow based on CRISPR/Cas9-mediated targeted mutagenesis in zebrafish. Nat Protoc. 2016;11(12):2357–75.

53. Liu JX, Zhang D, Xie X, Ouyang G, Liu X, Sun Y, et al. Eaf1 and Eaf2 negatively regulate canonical Wnt/beta-catenin signaling. Development. 2013;140(5):1067–78.

54. Liu JX, Xu QH, Li S, Yu X, Liu W, Ouyang G, et al. Transcriptional factors Eaf1/2 inhibit endoderm and mesoderm formation via suppressing TGF-beta signaling. Biochim Biophys Acta Gene Regul Mech. 2017;1860(10):1103–16.

55. Cui B, Ren L, Xu QH, Yin LY, Zhou XY, Liu JX. Silver_ nanoparticles inhibited erythrogenesis during zebrafish embryogenesis. Aquat Toxicol. 2016;177:295–305.

56. Zhou XY, Zhang T, Ren L, Wu JJ, Wang WM, Liu JX. Copper elevated embryonic hemoglobin through reactive oxygen species during zebrafish erythrogenesis. Aquat Toxicol. 2016;175:1–11.

57. Xu QH, Guan PP, Zhang T, Lu C, Li GL, Liu JX. Silver nanoparticles impair zebrafish skeletal and cardiac myofibrillogenesis and sarcomere formation. Aquat Toxicol. 2018;200:102–13.

